# Structural basis of quinone-sensing by the MarR-type repressor MhqR in *Staphylococcus aureus*

**DOI:** 10.1101/2025.10.30.685541

**Authors:** Thao Thi-Phuong Nguyen, Paul Weiland, Vu Van Loi, Stephan Kiontke, Fabiana Burchert, Victor Zegarra, Antonia Kern, Verena Nadin Fritsch, Daniil Baranov, Agnieszka K. Bronowska, Gert Bange, Haike Antelmann

## Abstract

The MarR-family regulator MhqR of *Staphylococcus aureus* (*Sa*MhqR) was previously characterized as quinone-sensing repressor of the *mhqRED* operon. Here, we resolved the crystal structures of apo-*Sa*MhqR and the 2-methylbenzoquinone (MBQ)-bound *Sa*MhqR complex. AlphaFold3 modelling was used to predict the structure of the *Sa*MhqR in complex with its operator DNA. In the DNA-bound *Sa*MhqR state, S65 and S66 of an allosteric α3-α4 loop adapted a helically wound conformation to elongate helix α4 for optimal DNA binding. Key residues for MBQ interaction were identified as F11, F39, E43, and H111, forming the MBQ-binding pocket. MBQ binding prevented the formation of the extended helix α4 in the allosteric loop, leading to steric clashes with the DNA. Molecular dynamics (MD) simulations revealed an increased intrinsic dynamics within the allosteric loop and the ß1/ß2-wing regions after MBQ binding, to prevent DNA binding. Using mutational analyses, we validated that F11, F39, and H111 are required for quinone sensing *in vivo,* whereas S65 and S66 of the allosteric loop and D88, K89, V91 and Y92 of the ß1/ß2-wing are essential for DNA binding *in vitro* and *in vivo*. In conclusion, our structure-guided modelling and mutational analyses identified a quinone-binding pocket of *Sa*MhqR and the mechanism of *Sa*MhqR inactivation, which involves local structural rearrangements of an allosteric loop and a high intrinsic dynamics to prevent DNA interactions. Our results provide novel insights into the redox-mechanism of the conserved *Sa*MhqR repressor, that functions as an important determinant of quinone and antimicrobial resistance in *S. aureus*.

**IMPORTANCE:** *S. aureus* is a major human pathogen, which can cause life-threatening infections in humans. However, treatment options are limited due to the prevalence of antimicrobial resistant isolates in the hospital and the community. The MarR-type repressor *Sa*MhqR was described to control resistance towards quinones and quinone-like antimicrobials. However, the redox-regulatory mechanism of *Sa*MhqR by quinones was unknown. In this work, we explored the DNA-binding and quinone-sensing mechanism of *Sa*MhqR and identified a quinone-binding pocket and an allosteric loop, which facilitates DNA binding activity via a helical wound conformation and adapts an unstructured coiled conformation upon quinone binding to inhibit DNA binding. A similar mechanism has been recently discovered for regulation of uric acid resistance by UrtR family repressors (1). Our results contribute to a better understanding of antimicrobial resistance regulation, which can be exploited for future drug-design to eradicate multidrug-resistant *S. aureus*.

## Introduction

*Staphylococcus aureus* is an opportunistic pathogen, which colonizes the skin and the nose of 20-60% of the healthy human population without causing symptoms of infections (2). However, under the hospital settings and in immunocompromised patients, *S. aureus* can develop to a major pathogen leading to local skin and soft tissue infections, but also to life-threatening diseases, such as bacteremia, osteomyelitis, pneumonia or catheter-associated endocarditis (2–5). The limited availability of vaccines and treatment failures due to antibiotic resistant and persistent strains poses a major health burden and high mortality rates of *S. aureus* bloodstream infections (6–8). Thus, there is an urgent need to elucidate the adaptation mechanisms of this important human pathogen under infection conditions with the aim of identifying novel drug targets and treatment options to eradicate multidrug-resistant *S. aureus* strains.

During infection, *S. aureus* is phagocytosed by human macrophages and neutrophils, which produce a range of highly microbiocidal compounds, including reactive oxygen species (ROS) and (pseudo-)hypohalous acids (HOCl, HOSCN) to kill the invading pathogen as first line of our innate immune defense (9, 10). In eukaryotic and prokaryotic cells, ROS can react with many cellular macromolecules, such as carbohydrates, unsaturated fatty acids, amino acids or other metabolites to generate secondary reactive species, such as reactive electrophile species (RES) (10–13). Bacterial pathogens might encounter RES, such as quinones, aldehydes, methylglyoxal or epoxides during respiration, cellular metabolism or host-pathogen interactions (10). In addition, many antimicrobial and toxic xenobiotic compounds are produced during microbial interactions and could be potent sources of ROS and RES for bacteria.

We have previously investigated the regulatory mechanisms of *S. aureus* under quinone stress, including 2-methylhydroquinone (MHQ) and the 1,4-naphthoquinone lapachol (10, 14–16). Quinones have two modes of actions, in the oxidative mode semiquinone anions lead to superoxide production, and as electrophiles protein thiols are irreversibly alkylated and aggregated by quinones (17–21). While benzoquinones act via both the oxidative and alkylation modes (20), lapachol was shown to exert its toxicity mainly via ROS formation (14). Thus, in solution MHQ is constantly autoxidized to form 2-methylbenzoquinone (MBQ), resulting in a color change from colorless to yellow-orange.

The response to quinones in *S. aureus* is governed by the two MarR-type repressors, MhqR and QsrR, which sense and respond to quinones and together control quinone detoxification pathways, including the phospholipase/carboxylesterase MhqD, putative ring-cleavage dioxygenases (MhqE, CatE, CatE2) and quinone reductases (AzoR1, YodC) (15, 16, 22). The MhqR and QsrR regulons conferred independent protection against MHQ and antimicrobial compounds, such as pyocyanin, ciprofloxacin, norfloxacin and rifampicin in *S. aureus* (15, 16, 23). The QsrR regulon was also induced by strong oxidants, such as diamide, HOCl, allicin and AGXX stress in *S. aureus* (14, 16, 24–27). We have shown that QsrR functions as two-Cys-type redox-sensing regulator, which senses quinones and oxidants via intersubunit disulfide formation between the redox-sensitive Cys4 and either Cys29’ or Cys32’ of the opposing subunit (15). Additionally, a one-Cys-type QsrR variant was also shown to sense quinones by thiol-*S*-alkylation of Cys4 *in vitro* (22).

However, the MhqR repressor does not possess a conserved Cys residue and its sensing mechanism is unknown (16). Many MarR-type repressors sense specific ligands, such as antibiotics, phenolic compounds, organic hydroperoxide, uric acid or other metabolites, which bind to a conserved ligand-binding pocket, resulting in structural and conformational alterations, incompatible with DNA binding (1, 28–32). In this work, we have investigated the structural changes of the DNA-bound MhqR repressor of *S. aureus* (*Sa*MhqR) upon MHQ binding. We used mutational, biochemical and crystallographic analyses, along with AlphaFold3 modelling and molecular dynamics (MD) simulations, to identify the MBQ-binding pocket and to elucidate the structural and dynamic changes of an α3-α4 allosteric loop and within the wHTH motif upon quinone binding, resulting in dissociation of the *Sa*MhqR repressor from the promoter DNA.

## Results

### High-resolution structure of *Sa*MhqR reveals a stable dimer resembling typical MarR-type repressors

We have previously characterized the MhqR repressor of *S. aureus*, which senses quinones and controls the *mhqRED* operon, conferring resistance towards MHQ stress and quinone-like antimicrobials (16). *Sa*MhqR consists of 144 amino acids with a molecular weight of 16.5 kDa. To elucidate the three-dimensional protein structure of *Sa*MhqR, we employed X-ray crystallography, solving its structure to a resolution of 1.55 Å **(Table S1, PDB: 9QDR)**. The crystal structure reveals a homodimer with a fold typical of MarR-family regulators (28–30) **(Fig. 1A)**. Each subunit of the *Sa*MhqR dimer comprises six α-helices, with helices α4 and α5 flanking a β-hairpin structure (ß1/ß2-wing). The DNA binding domain is composed of the helices α2, α3 and α4 and the ß1/ß2-wing, forming the characteristic winged helix-turn-helix (wHTH) motif **(Fig. 1A)**.

**Figure 1.**
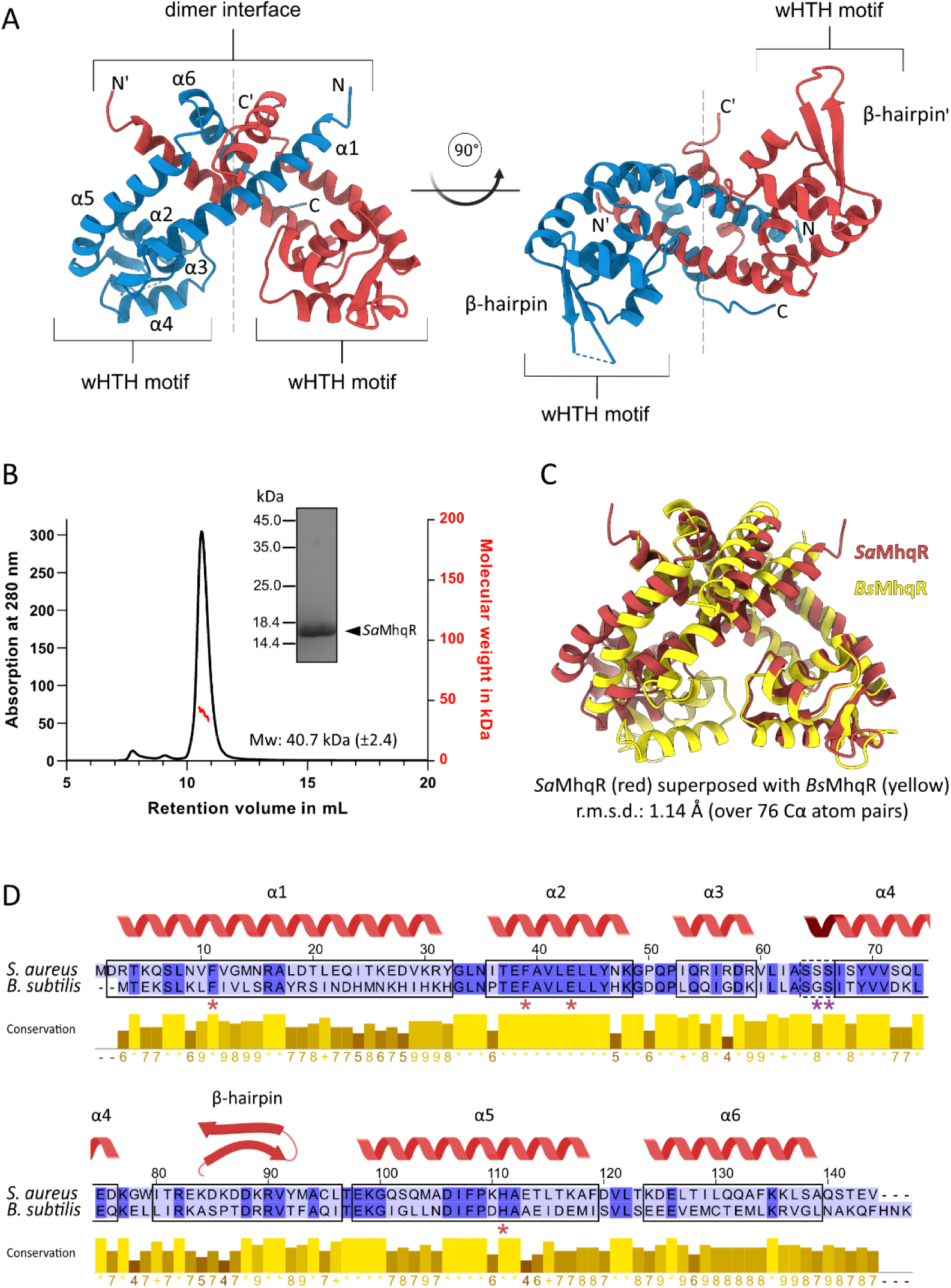
**Crystal structure and dimer formation of the quinone-sensing *S. aureus* MarR-type repressor *Sa*MhqR**. **A)** The crystal structure of *Sa*MhqR reveals a classical MarR-type protein fold with two monomers (red and blue) in the dimeric structure. The winged helix-turn-helix motifs (wHTH) are formed by α-helices α2, α3, and α4 and the ß-hairpin, facing outwards. The α-helices α1, α5, and α6 contribute to the formation of the dimer interface. **B)** Using analytical SEC-MALS, the molecular weight of purified *Sa*MhqR was determined as 40.7 kDa (±2.4) confirming that *Sa*MhqR is present as dimer in solution. **C)** Superposing the structure of the *Sa*MhqR with the predicted structure of *Bs*MhqR reveals a high structural similarity. **D)** Sequence alignment of *Sa*MhqR and *Bs*MhqR using ClustalΩ2 (http://www.clustal.org/omega/) reveals an overall 39.4% sequence identity, and a very high conservation of the α-helix α2. The quinone-binding residues (Phe11, Phe39A, Glu43 and His111) and the DNA-binding residues Ser65 and Ser66 in the α3-α4 allosteric loop, which are essential for the elongation of α4 upon DNA binding of *Sa*MhqR, are marked by red and violet stars, respectively.

The two *Sa*MhqR subunits form a robust dimer. Helices α1, α5, and α6 of each subunit contribute to a large buried interface, covering approximately 2890 Å² with an estimated solvation energy of around 111 kcal/mol for each monomer, making it an energetically favorable oligomerization state **(Fig. S1A)**. The dimeric state of *Sa*MhqR in solution was confirmed by size-exclusion chromatography coupled with multi-angle light scattering (SEC-MALS) indicating a molecular weight of 40.7 ± 2.4 kDa **(Fig. 1B)**. Notably, the artificial C-terminal hexahistidine tag appears to aid crystallization by interacting with the dimer interface of *Sa*MhqR symmetry mates in neighboring asymmetric units, and is not depicted in the main structural figures.

We have previously studied also the role of the MhqR homolog of *B. subtilis* (*Bs*MhqR), revealing similar functions of its regulon members to mediate quinone resistance (33). To investigate the degree of conservation between *Sa*MhqR and *Bs*MhqR, we predicted the structure of *Bs*MhqR using AlphaFold3 (34). The resulting model showed high confidence **(Fig. S2A)** and strong structural similarity to *Sa*MhqR **(Fig. 1C; Fig. S2B)**. Structural superposition of *Sa*MhqR with the *Bs*MhqR model revealed a root mean square deviation (RMSD) of 1.17 Å over 110 paired Cα atoms and a sequence identity of approximately 39.4% **(Fig. 1C)**. Interestingly, sequence and structural alignments indicate that α-helix 2 is highly conserved between both MhqR homologs **(Fig. 1C, D)**.

Taken together, our crystal structure of *Sa*MhqR confirms its classification as a MarR-type protein, characterized by a typical dimeric structure and a wHTH motif, a hallmark of DNA-binding transcriptional regulators (28–30). *Sa*MhqR forms an energetically strong homodimer, as evidenced by its large buried interface and favorable solvation energy. Furthermore, it exhibits both structural and sequence similarity to *Bs*MhqR, underscoring its conserved function in the quinone stress response and similar architecture across Firmicutes.

### MhqR binds to its operator DNA by stabilizing and elongating α-helix 4

To investigate the conformational changes of *Sa*MhqR, we utilized AlphaFold3 modelling (34) to predict the structure of the *Sa*MhqR dimer bound to its double-stranded operator DNA, a 9-9 bp palindromic sequence (TATCTCGAA-ATCGAAATA) as identified in the *mhqRED* promoter previously **(Fig. 2A)** (16). The prediction of the *Sa*MhqR structure in complex with DNA yielded high pLDDT scores, indicating strong confidence in the modeled structure **(Fig. S3A)**. To further support our predicted model and the specific recognition of the palindromic sequence by the *Sa*MhqR dimer, we modelled DNA binding to a scrambled operator sequence with the same ATCG content as the native operator **(Fig. S3B)**. As expected, the model of SaMhqR in complex with the scrambled (non-cognate) operator resulted in a low pLDDT score, indicating a low confidence **(Fig. S3B)**. Thus, the model of *Sa*MhqR in complex with its cognate operator supports the specificity of DNA binding **(Fig. S3A)**. Using DNAproDB (35), we analyzed and illustrated the amino acid residues involved in the predicted DNA interactions of the *Sa*MhqR DNA complex and identified α-helix 4 as a key contributor to both base-pair recognition and sugar-phosphate backbone interactions within the major groove of the dsDNA helix **(Fig. 2B)**.

**Figure 2.**
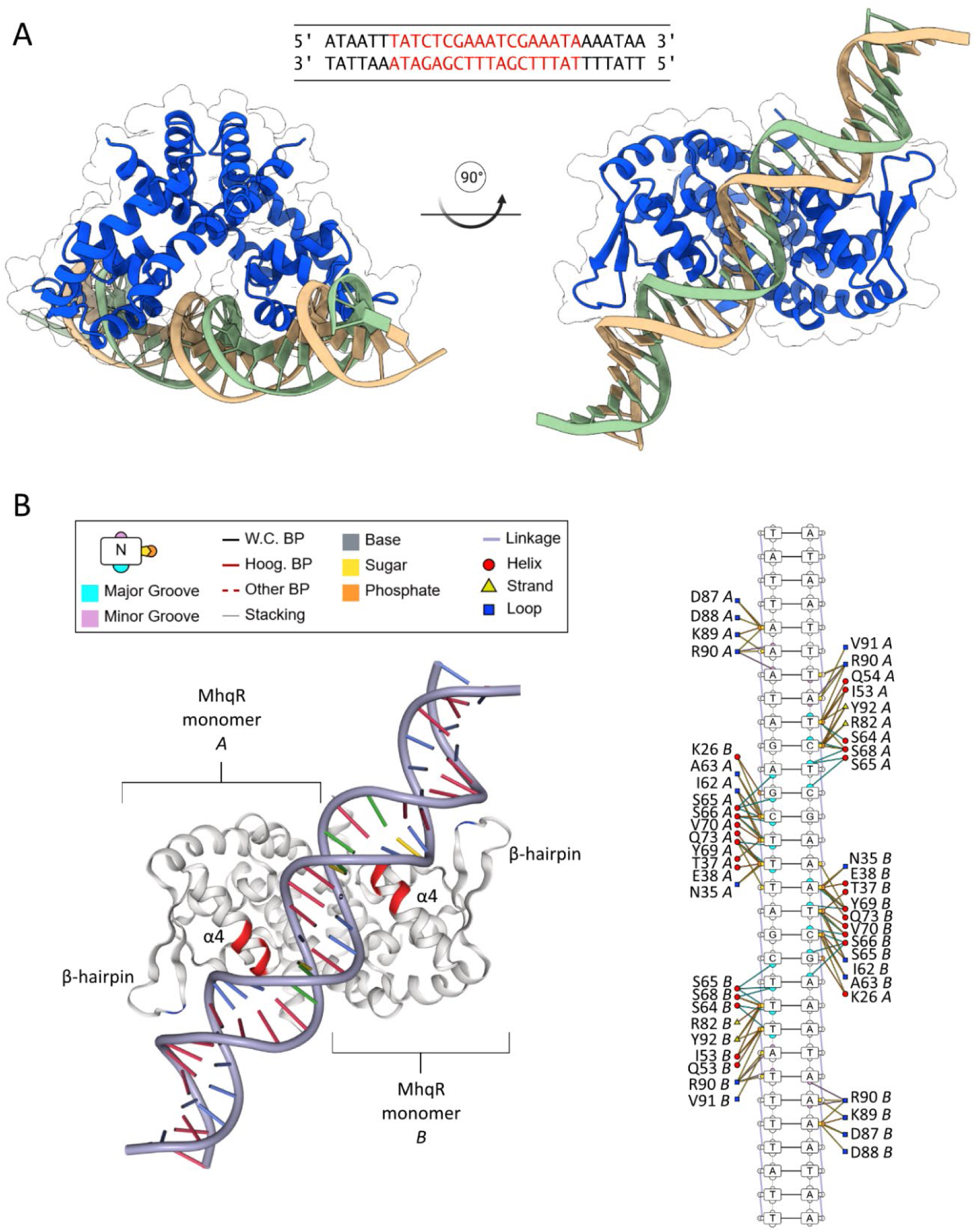
Predicted structure of *Sa*MhqR in complex with its operator DNA upstream of the *mhqRED* operon. **A)** The predicted DNA-protein complex of the *Sa*MhqR dimer (blue) bound to its operator DNA sequence (beige and green) was modelled with AlphaFold3 (34). The sequence of the palindromic *Sa*MhqR operator is shown at the top (red). **B**) Amino acid residues of each monomer contributing to the recognition of specific nucleotide bases, phosphate backbone, and sugars of the operator sequence are listed as well as their corresponding secondary structures indicating that mainly α-helix α4 and partly the β-hairpin are involved in DNA recognition.

Residues responsible for base-pair recognition in α-helix 4 include serine S64, S65, S66, and S68, as well as tyrosine Y69. Interestingly, in the DNA-bound conformation, α-helix 4 exhibits an extended structure compared to its apo state **(Fig. 2B)**. Notably, the serine residues S65 and S66 of the α3-α4 loop transition from being unstructured in the apo state to a helical wound conformation contributing to the elongated α-helix 4 during DNA binding, underscoring its pivotal role in the interaction **(Fig. 1A**, **Fig. 2A, B)**. To investigate the crucial roles of S65 and S66 of the α3-α4 loop in DNA-binding, we modelled the structure of the *Sa*MhqR S65A,S66A double mutant in complex with the cognate operator DNA **(Fig. S3C)**. The *Sa*MhqR S65A,S66A variant was unable to recognize with its wHTH motifs the adjacent major and minor groves of the DNA double helix, as indicated by low pLDDT score and therefore low confidence of the model (**Fig. S3C)**. Thus, our *in silico* predictions strongly support that the S65 and S66 residues of the α3-α4 loop are essential for DNA binding of the *Sa*MhqR dimer.

Beyond helix α4, additional interactions seem to stabilize the DNA-bound conformation of *Sa*MhqR. Glutamine Q54 from loop 3 engages with the major groove of the dsDNA, while arginine R90 from the β-hairpin loop forms stabilizing interactions with adenine and thymine in the minor groove **(Fig. 2B)**.

Other regions, including helices α2 and α3 as well as the β1/ß2-wing, contribute to DNA coordination via non-specific interactions with the sugar-phosphate backbone **(Fig. 2B)**. Notably, all key residues involved in DNA binding are conserved between *Bs*MhqR and *Sa*MhqR.

Altogether, our *in silico* analyses support that *Sa*MhqR binds specifically to its operator DNA through specific and non-specific interactions, with α-helix 4 playing a central role in base-pair recognition and sugar-phosphate backbone interactions. The extension of α-helix 4 in the DNA-bound state, particularly the involvement of S65 and S66 in the α3-α4 loop, underscores its pivotal role in stabilizing the complex.

### MBQ binding allosterically prevents DNA binding of *Sa*MhqR

We previously hypothesized that *Sa*MhqR senses quinones via direct binding to a specific ligand pocket (16). Thus, we used isothermal titration calorimetry (ITC) to investigate the binding of *Sa*MhqR to both reduced MHQ and oxidized MBQ, since MHQ is known to undergo redox-cycling **(Fig. 3A, B)**. The ITC results showed that *Sa*MhqR binds to MHQ with a dissociation constant (K_D_) of ∼ 6 ± 0.5 µM and slightly stronger to MBQ with a K_D_ of ∼3 ± 0.6 µM **(Fig. 3B)**.

**Figure 3.**
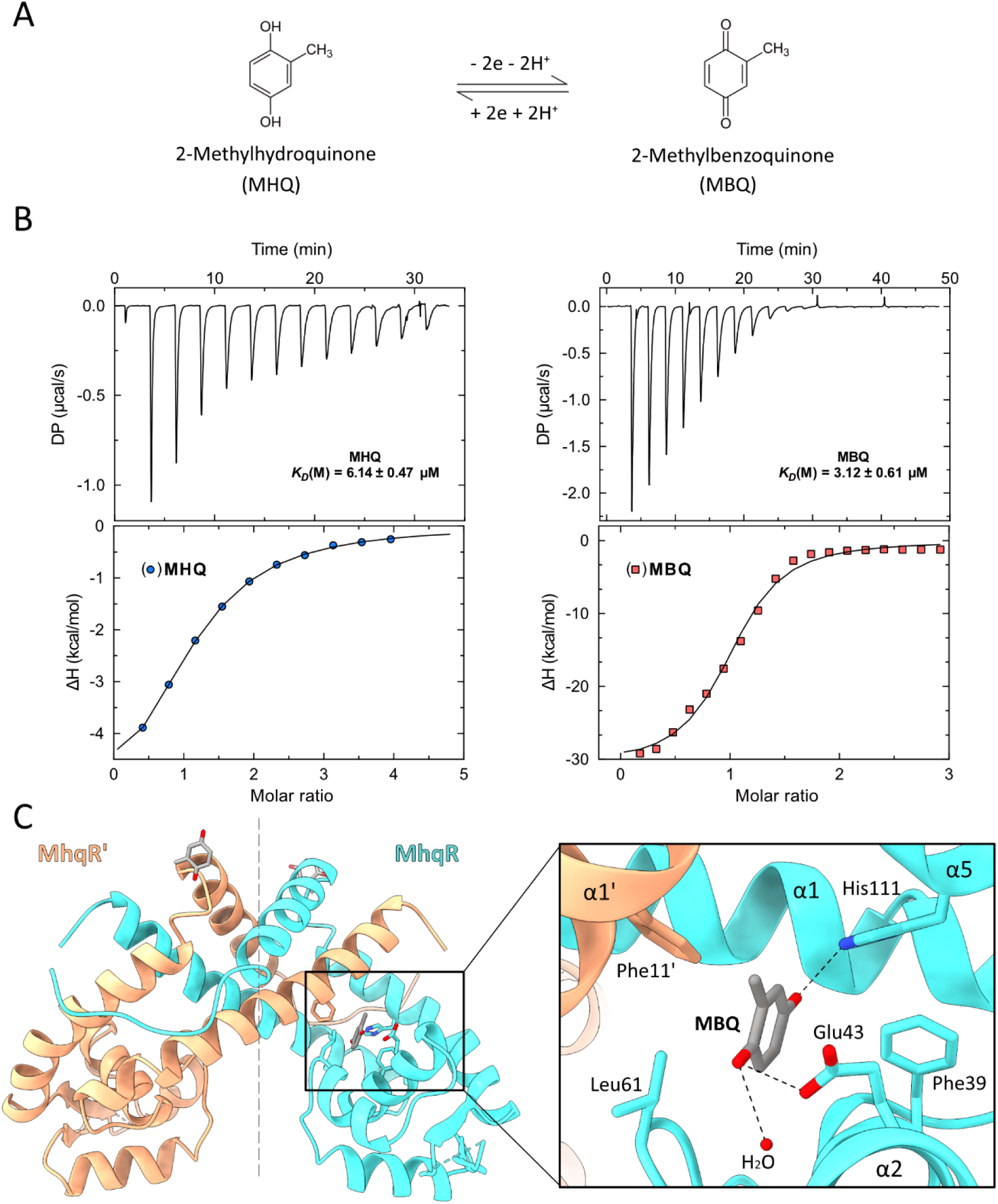
Binding of MBQ to the ligand pocket of *Sa*MhqR, **A)** The chemical structures and redox cycles of reduced 2-methylhydroquinone (MHQ) and oxidized 2-methylbenzoquinone (MBQ). **B)** The affinity of *Sa*MhqR to MHQ and MBQ was determined by quantifying the K_D_ via ITC assays and was calculated as ∼ 6 µM for MHQ and ∼3 µM for MBQ. The ITC assays were performed in 2 replicates and one representative experiment is shown. **C)** The crystal structure of the *Sa*MhqR-MBQ complex reveals that the MBQ binding site is formed by the α-helices α2 and α5 of one subunit (cyan) and α-helices α1 of the opposing subunit (orange). A close-up view reveals that the amino acids Phe11, Phe39, Glu43 and His111 are involved in the formation of the MBQ binding pocket, with Glu43 and His111 close enough for the formation of hydrogen bonds to MBQ (dashed black lines).

The quinone-binding pocket of MarR-type repressors is highly conserved, though it has not been identified in *Sa*MhqR. To address this, we determined the crystal structure of the *Sa*MhqR dimer in complex with MHQ **(Fig. 3C; PDB: 9SMZ)**. Since the obtained crystals were yellow, similar to the oxidation product MBQ in solution, we assumed that MBQ was bound as a ligand in the binding pocket of *Sa*MhqR. After phase determination and initial refinement, positive Fo-Fc electron density was clearly observed only in the binding pocket of molecule A. Refinement including MBQ as a ligand with an overall occupancy of 80% (q = 0.8) in two orientations (q_1_ = 0.5; q_2_ = 0.3) resulted in an unambiguous 2Fo-Fc electron density **(Fig. S4A)**. To further verify the presence of MBQ, a final polder omit map was calculated, excluding bulk solvent around the omitted region (36). The resulting map clearly revealed the omitted atoms, thereby confirming the presence of MBQ in the binding pocket **(Fig. S4B)**. The MBQ binding pocket is located between α-helices α2 and α5 of one subunit, and α-helix α1 of the opposing subunit. Key amino acids involved in the formation of the MBQ binding pocket include H111 of α5, F39 and E43 of α2, and F11’ of α1’ of the opposing subunit of *Sa*MhqR. The two carbonyl groups of MBQ were shown to be coordinated by the conserved H111 residue, E43 and a water molecule **(Fig. 3C)**. The distance between the MBQ carbonyl oxygen to H111 and E43 was measured as 2.91 Å and 2.78 Å, respectively, indicating the formation of moderate hydrogen bonds with the ligand **(Fig. 3C)**.

We further compared the MBQ-binding pocket of SaMhqR with the ligand-binding sites of other MarR-family proteins. The MBQ ligand site was found at a similar position as the ligand pockets of the MarR-type regulators Rv2887 of *Mycobacterium tuberculosis* (PDB: 5X80) (37, 38), ST1710 of *Sulfurisphaera tokodaii* (PDB: 3GF2) (39), and AbsC of *Streptomyces coelicolor* (PDB: 3ZMD) (40), all bound to salicylic acid **(Fig. S5A, C)**. Additionally, we compared the MBQ-bound *Sa*MhqR structure with the structure of HucR from *Deinococcus radiodurans* bound to urate (PDB: 7XL9) and found the same binding pocket **(Fig. S5C)** (41, 42).

Sequence alignments of these MarR proteins revealed that most of the MBQ pocket residues of *Sa*MhqR are conserved in other ligand-binding sites, such as F39 in *Sa*MhqR, Rv2887 and ST1710, E43 in *Sa*MhqR and AsbC, and H111 in *Sa*MhqR, AsbC and HucR **(Fig. S5B, D)**. Other residues appear to be variable, suggesting that the binding sites have evolved different binding modes to accommodate a diverse range of ligands.

Interestingly, superposition of the DNA-bound and MBQ-bound *Sa*MhqR structures revealed that the α3-α4 loop intrudes into the ligand pocket of MBQ-bound *Sa*MhqR, resulting in a steric clash with the DNA backbone and dissociation of SaMhqR from the operator **(Fig. 4A)**. This loop plays an important role for the interaction of *Sa*MhqR with DNA, as it needs to remain flexible to allow the stabilization and elongation of α-helix α4 during DNA binding **(Fig. 4A)**. The binding of MHQ or MBQ may thus prevent this conformational change and stabilize the extruded form of this loop, preventing α-helix α4 from adopting the elongated form necessary for proper DNA binding **(Fig. 4A)**.

**Figure 4.**
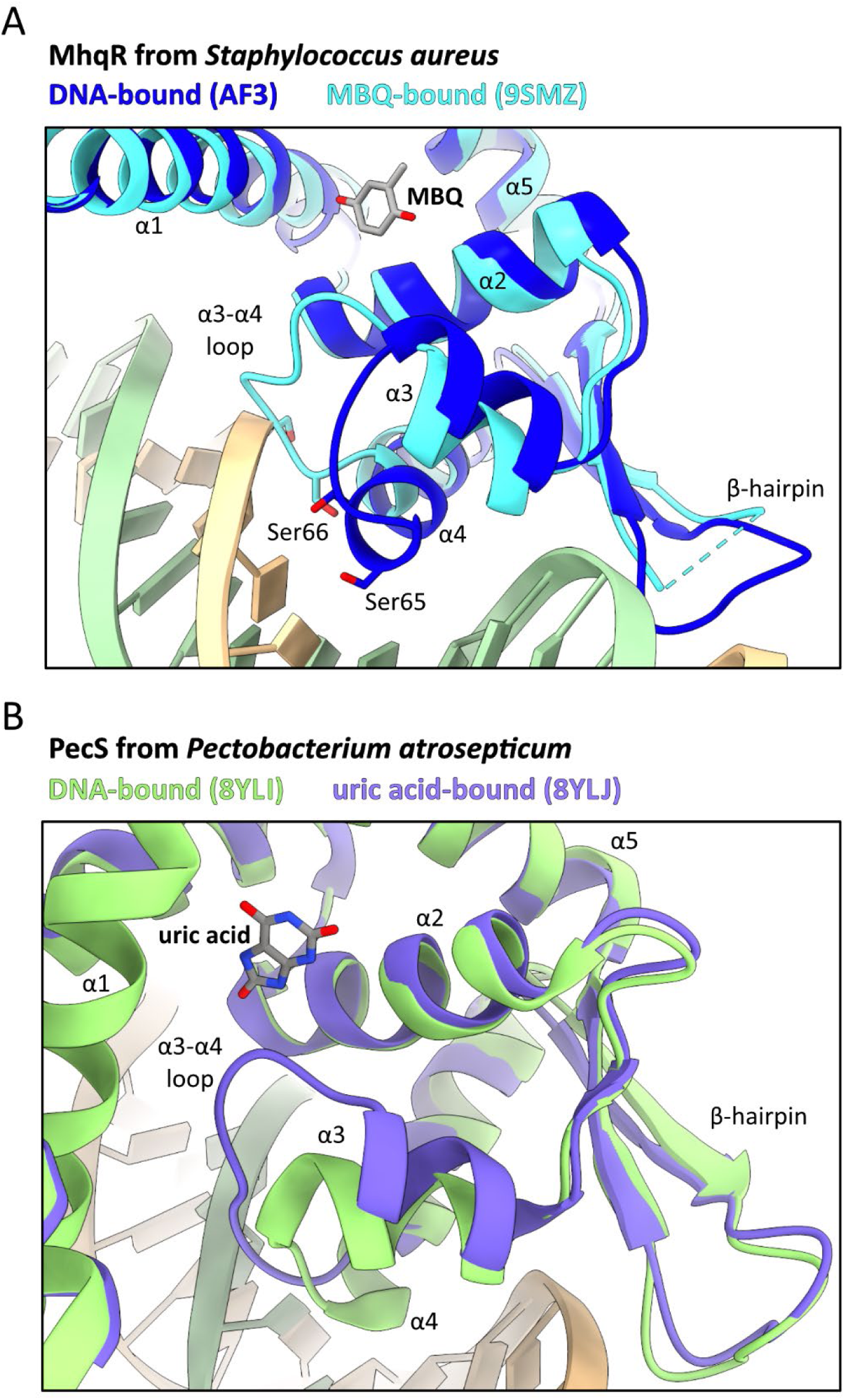
Conformational changes of the *Sa*MhqR-DNA complex. **A)** Superposing the predicted DNA-*Sa*MhqR complex (blue) and the experimentally determined *Sa*MhqR-MBQ complex (cyan) shows that Ser65 and Ser66, which are essential for DNA binding, elongate the length of α-helix α4 and therefore shortening the α3-α4 loop to facilitate DNA binding. Upon MBQ binding, the allosteric α3-α4 loop adopts a coiled conformation, disrupting DNA binding of *Sa*MhqR. **B)** Close-up views illustrate the conformational changes in α-helix α4 of PecS from *Pectobacterium atrosepticum* upon uric acid binding as determined previously (1). These changes extend to the α3-α4 loop when comparing the DNA-bound state (8YLI, green) with the uric acid-bound (8YLJ, purple) states.

These findings closely resemble the structural changes observed upon uric acid binding in UrtR family proteins, such as PecS, MftR, and HucR, that interfere with DNA binding (1). In UrtR proteins and *Sa*MhqR, an elongated and stabilized α4-helix is critical for DNA binding. Similarly, the adjacent α3-α4 loop is involved in the formation of the ligand-binding pocket and influences proper DNA binding upon uric acid binding **(Fig. 4B)** (1).

In conclusion, we have shown that *Sa*MhqR binds MHQ with a K_D_ of ∼6 ± 0.5 µM, and MBQ with a K_D_ of ∼3 ± 0.6 µM, as demonstrated by ITC. Crystal structure analysis of the *Sa*MhqR-MBQ complex identified a binding pocket involving the conserved ligand-binding residues F11, F39, E43 and H111. Comparisons with other MarR-type proteins confirmed the conservation of the ligand pocket across ligand-binding MarR-type regulators (28). MBQ binding stabilizes the α3-α4 loop, critical for DNA interaction, resulting in a steric clash with the DNA and dissociation from its operator. Thus, the regulatory mechanism of quinone sensing by SaMhqR was shown to be identical to that of uric acid-sensing by PecS from *Pectobacterium atrosepticum* **(Fig. S4B)** (1).

### Molecular dynamic (MD) simulations of DNA- and MHQ-bound *Sa*MhqR

We further used atomistic MD simulation to investigate the structural changes and to predict protein intermediate conformational states of the DNA-bound *Sa*MhqR complex upon MHQ binding. Analysis of the MD simulation trajectories showed that the key structural difference of DNA-bound to MHQ-bound *Sa*MhqR occurs in the α3-α4 allosteric loop region, spanning residues 58-68, and in the ß1/ß2-wing region (residues 80-96) **(Fig. S6, S7 and S8)**. The quantification of the secondary structure content over MD simulation time of 100 ns was performed using the Define Secondary Structure of Proteins (DSSP) algorithm for DNA- and MHQ-bound *Sa*MhqR, revealing major structural changes in the region between Leu61 and Ser66 of the allosteric loop upon MHQ-binding **(Fig. S7A, B)**. We observed that the conformation of this allosteric loop in the MHQ-bound *Sa*MhqR complex would disfavor DNA binding due to a steric clash **(Fig. S6)**. The overall stability of apo *Sa*MhqR and the DNA- and MHQ-bound *Sa*MhqR complexes was evaluated by root mean square deviation (RMSD) **(Fig. S8A-C)**, and local flexibility was assessed via per-residue root mean square fluctuation (RMSF) **(Fig. S8D-F)**. The RMSF plots showed high fluctuations in the ß1/ß2-wing region (residues 80-96), which contact the minor groove of the DNA operator **(Fig. S8D-F)**.

Moreover, we also sampled the motions associated with the transition from the DNA-bound to the MHQ-bound *Sa*MhqR protein using coarse-grained Elastic network-driven Brownian Dynamics IMportance Sampling (eBDIMS), followed by reverse-mapping to all-atom resolution **(Fig. 5A-C)**. The simulation data revealed that binding of either DNA, or MHQ affects the *Sa*MhqR intrinsic dynamics overall **(Fig. 5A-C)**, but in different ways. Principal component analysis (PCA) indicates that, relatively to the DNA-bound state, MHQ binding increased the intrinsic dynamics of the α3-α4 allosteric loop region (residues 58-68) and the ß1/ß2-wing (residues 80-96) of the wHTH motif, which are both essential for DNA-binding **(Fig. 5A-C)**. This is likely to prevent DNA binding upon MHQ binding due to entropic penalty in the ß1/ß2-wing region. Thus, our simulation data enabled to describe the “dynamic allostery” between the DNA-bound and MHQ-bound states of *Sa*MhqR, showing an increased intrinsic dynamic in the ß1/ß2-wing upon MHQ-binding.

**Figure 5.**
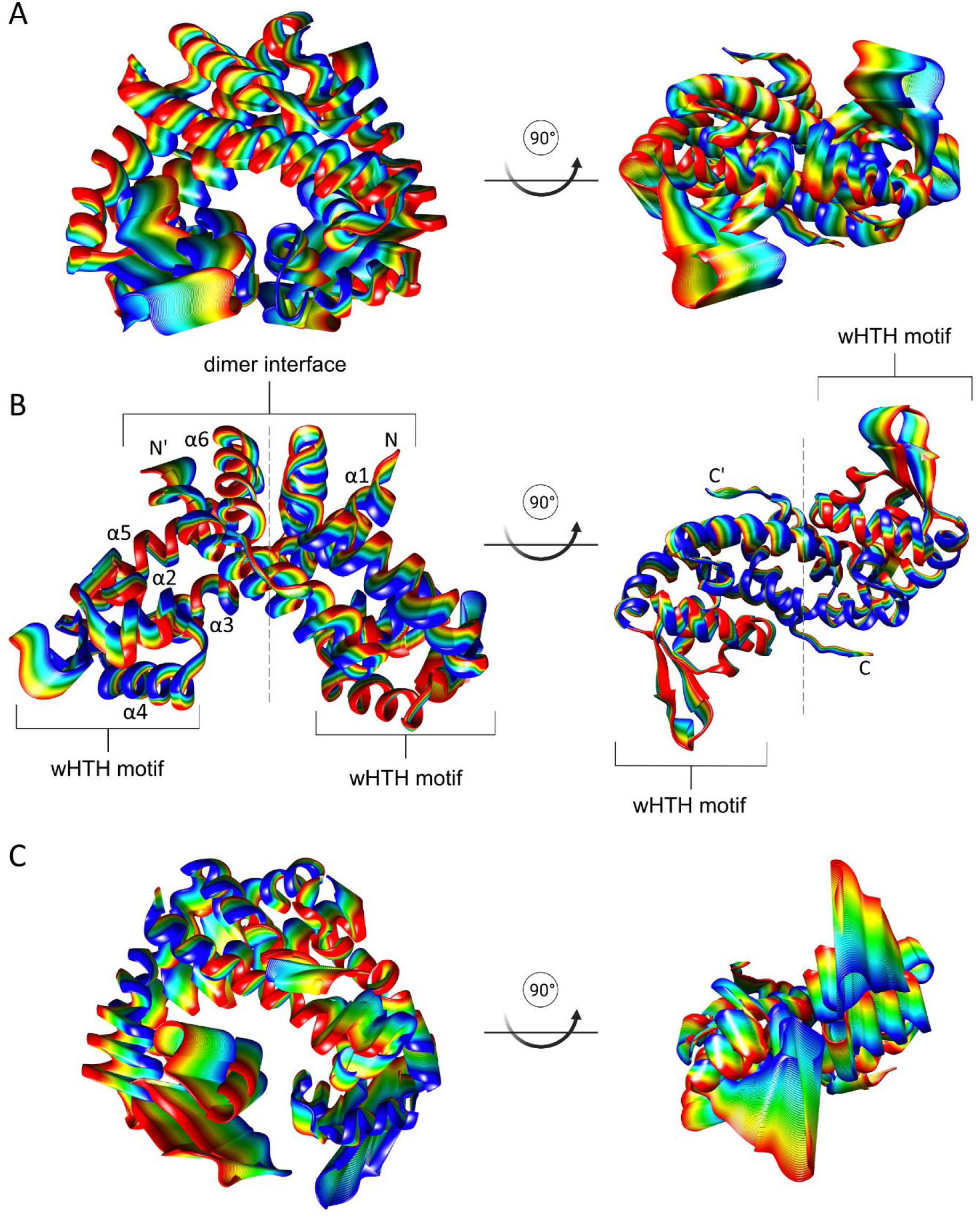
Ribbon plots of the first principal component (PC1), obtained from molecular dynamics simulations. The PCA analysis was performed after coarse-grained simulation for the apo *Sa*MhqR dimer **(A)**, DNA-bound *Sa*MhqR dimer **(B)**, and MHQ-bound *Sa*MhqR dimer **(C)** (left - side view, right - bottom view). Colors (from blue to red) show the progression between the extremes, and the width of the ribbon is proportional to the mobility.

### Effect of amino acid substitutions of *Sa*MhqR on quinone sensing and DNA binding activity *in vitro*

To investigate the significance of amino acids involved in the MHQ/MBQ binding pocket, which was identified in the crystal structure of the MBQ-bound *Sa*MhqR complex, we individually mutated the pocket residues F11, F39, E43 and H111 to alanine in *Sa*MhqR. In addition, we constructed the *Sa*MhqR E43A,H111A allele, since both residues were engaged in specific interactions with the carbonyl groups of the quinone **(Fig. 3C)**. We further aimed to validate the role of predicted DNA-binding residues of *Sa*MhqR by construction of various *Sa*MhqR variants with mutations in the α3-α4 allosteric loop (A63E, S64A, S65A, S66A), in the helices α2/ α3 (E38A, Q54A) and α4 (S68A, Y69A, V70A, Q73A) and in the ß1/ß2-wing (D88A, K89A, R90A, V91A, Y92A).

To confirm that all *Sa*MhqR variants were stable in their protein secondary structures, we analyzed the far-UV circular dichroism (CD) spectra of the variants in comparison to the *Sa*MhqR WT protein. Overall, the CD spectra of the 20 *Sa*MhqR variants mutated in the ligand pocket and the predicted DNA-binding residues were similar to that of the WT protein, indicating no major structural changes in the content of α-helices and ß-sheets of the variants **(Fig. S9)**. Thus, all variants maintained their high α-helical content as typical for MarR-type proteins.

Next, the purified *Sa*MhqR mutants in the ligand pocket residues were used for ITC binding assays to MHQ and MBQ in comparison to the *Sa*MhqR WT protein. The thermodynamic values and dissociation constants for each variant are summarized in **Table S2**. The fitted ITC curves for these mutants are provided in **Fig. S10**. However, none of the single mutants in the binding pocket residues caused a drastic change in the binding behavior towards MHQ or MBQ **(Fig. S10; Table S2)**.

To further investigate whether the purified *Sa*MhqR variants are impaired in MHQ sensing or DNA-binding, we performed electrophoretic mobility shift assays (EMSAs) **(Fig. 6A)**. The EMSA results of the *Sa*MhqR variants with mutations in the ligand pocket showed that E43A bound with similar affinity to the *mhqRED* operator DNA, whereas the binding affinities of F11A, F39A, and H111A were lower as compared to the WT protein. Furthermore, the E43A, H111A double mutant was unable to bind to the DNA **(Fig. 6A)**. This may reflect the close association of the quinone-binding residues with the adjacent α3-α4 loop, which is important for DNA binding. In addition, the DNA binding assays with MHQ-treated *Sa*MhqR variants in the ligand sites revealed a similar response as observed for *Sa*MhqR WT protein, where inactivation of the repressor was observed only with 20-50 mM MHQ **(Fig. 6A)**. Overall, our ITC and EMSA results revealed that the *Sa*MhqR variants with alanine substitutions in the ligand-binding residues did not affect the MHQ/MBQ binding and sensing ability of *Sa*MhqR *in vitro*. However, since mutation of even two ligand sites in the E43A, H111A double mutant affected the DNA binding activity, it was not possible to investigate the role of combinatory mutations in MHQ/MBQ sensing *in vitro*.

**Figure 6.**
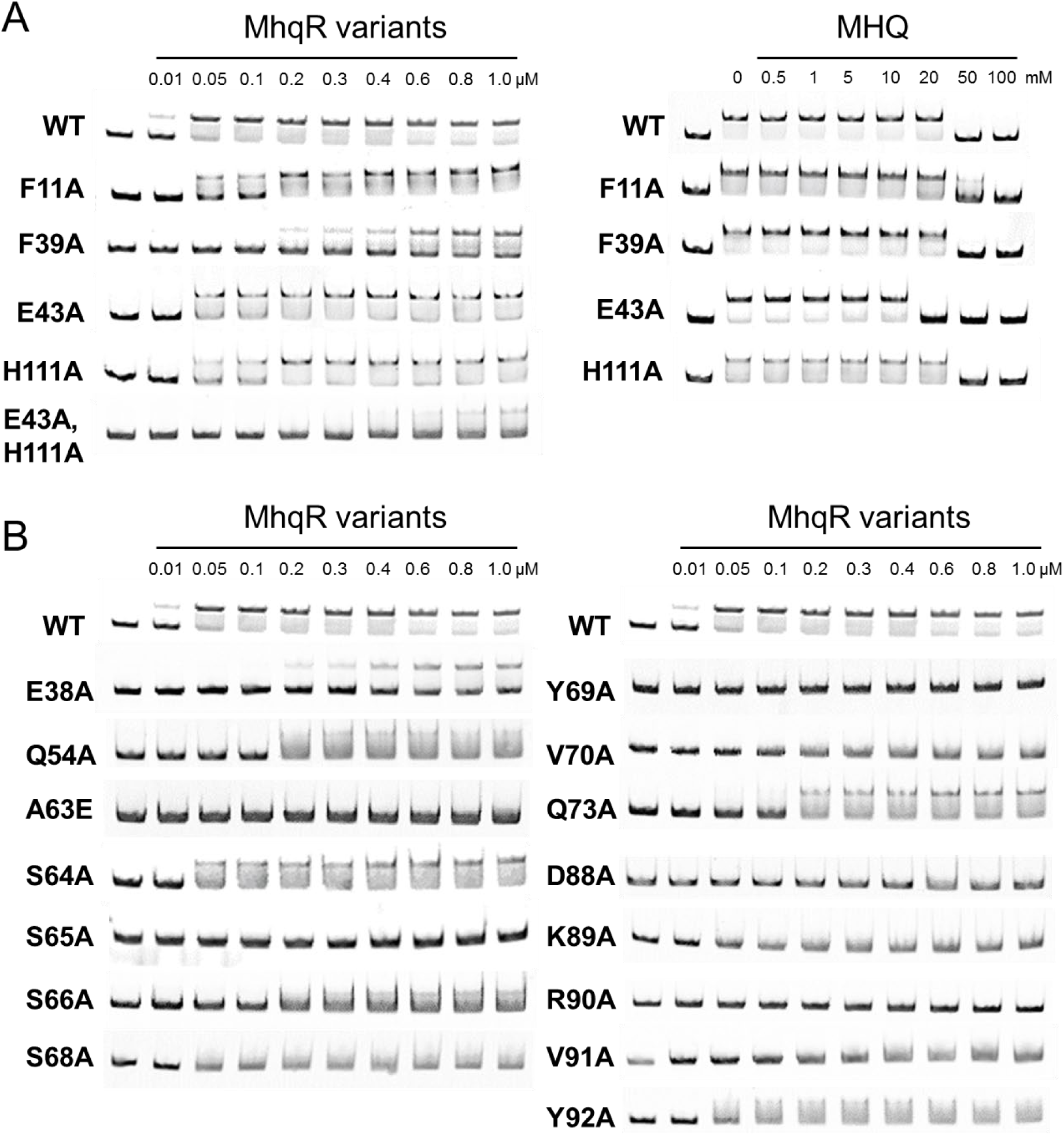
**Effect of amino acid substitutions on the DNA binding activity of purified *Sa*MhqR variants *in vitro***. EMSAs were performed to analyze the DNA-binding activity and MHQ response of *Sa*MhqR WT and the variants with mutation in **(A)** the ligand binding site (F11A, F39A, E43A, H111A and E43A,H111A) and **(B)** the predicted DNA-interaction sites (α2/α3: E38, Q54A), (α3-α4 loop: A63E, S64A, S65A, S66A), (α4: S68A, Y69A, V70A, Q73A) and the ß-hairpin (D88A, K89A, R90A, V91A, Y92A). *Sa*MhqR proteins of increasing concentrations (0.01-1.0 μM) were incubated with the *mhqRED* promoter DNA probe and subjected to EMSAs. **A)** Treatment with increasing concentrations of MHQ (50–100 mM) resulted in the dissociation of *Sa*MhqR WT and variants from the DNA probe. The experiments were performed in 3 replicates and one representative is shown.

Next, we used EMSAs to analyze the role of MhqR variants with mutations in the predicted DNA interaction sites for their impact on DNA binding activity. Consistent with the predictions of the DNA-bound complex, most selected variants with mutations in the α3-α4 allosteric loop (A63E, S65A, S66A), in the helices α2/ α3 (E38A, Q54A) and α4 (S68A, Y69A, V70A, Q73A) and in the ß1/ß2-wing (D88A, K89A, R90A, V91A, Y92A) were impaired in DNA binding *in vitro*, except for the S64A variant, which bound similar as the *Sa*MhqR WT protein to the promoter DNA **(Fig. 6B)**. These results validated that the critical serine residues S65 and S66 in the α3-α4 allosteric loop are important for DNA binding of *Sa*MhqR *in vitro*, confirming the structural model of the S65A,S66A variant, which was unable to bind with the wHTH motifs to the adjacent major grooves of the DNA double helix **(Fig. S3C)**. Moreover, the residues of the ß1/ß2-wing were also confirmed to be essential for DNA binding, which were predicted to make contacts with the minor grooves of the operator DNA. Overall, our mutational analyses in the predicted DNA-binding residues confirmed particularly the importance of residues S65 and S66 in α3-α4 allosteric loop and the ß1/ß2-wing for DNA binding of SaMhqR *in vitro*.

### The impact of *Sa*MhqR variants on MHQ sensing and DNA binding activity *in vivo*

Next, we confirmed in Northern blot analyses using a *mhqR*-specific probe that the *mhqR* variants encoded from plasmid pRB473 in *S. aureus* are transcriptionally expressed during growth with xylose **(Fig. S11)**. To assess the role of the residues involved in the MBQ/MHQ-binding pocket for *Sa*MhqR regulation *in vivo*, we used Northern blot and qRT-PCR analyses to study the transcription of the *mhqRED* operon in *S. aureus* strains expressing the variants in the pocket residues (F11A, F39A, E43A, H111A, and E43A,H111A) in response to 100 µM MHQ stress **(Fig. 7A)**. The results revealed that the F11A and F39A variants showed similar DNA binding repressor activity as the *mhqR*^+^ complemented strain under control conditions, indicating no effect on DNA-binding activity **(Fig. 7A)**. The basal transcription of the *mhqRED* operon was significantly higher in the E43A variant and lower in the H111A and E43A,H111A variants than the *mhqR*^+^ strain. Importantly, the variants F11A, F39A, H111A and E43A,H111A were significantly impaired in quinone-sensing as revealed by the lower response of the *mhqRED* operon to MHQ stress compared to the *mhqR*^+^ strain **(Fig. 7A)**. Especially, the H111A and E43A,H111A ligand mutants were completely impaired in MHQ sensing, since no transcript was visible under MHQ stress, supporting the crucial role of H111 for MHQ sensing *in vivo*. In summary, using transcriptional analysis we could validate the roles of the conserved residues F11, F39 and H111 of the MHQ/MBQ-binding pocket of *Sa*MhqR for MHQ sensing and regulation *in vivo*.

**Figure 7.**
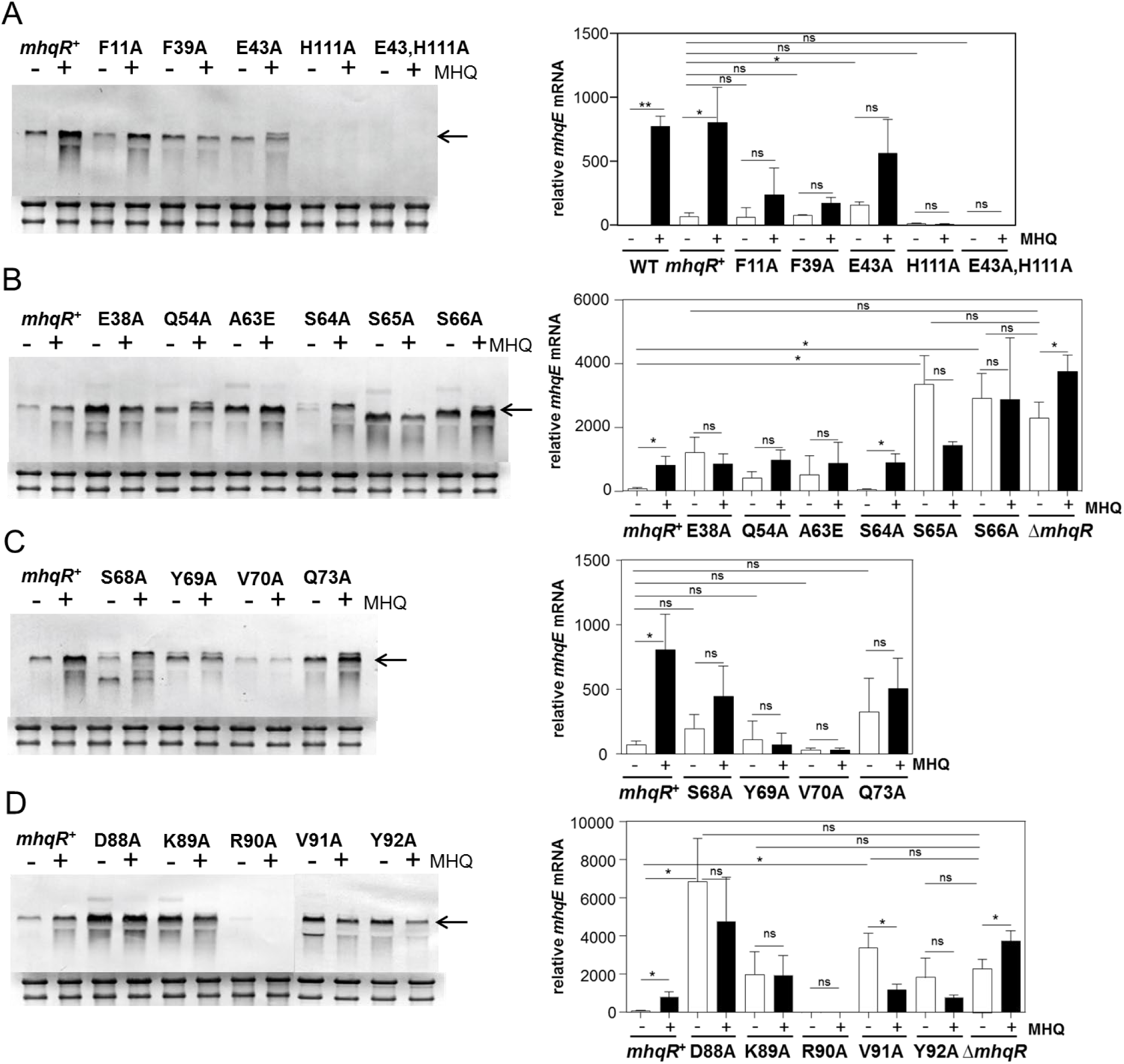
Transcription of the *mhqRED* operon in *S. aureus* strains expressing *Sa*MhqR variants. A-D) Northern blot and qRT-PCR analyses of *mhqRED* transcription were performed using RNA from *S. aureus* COL WT, the Δ*mhqR* mutant, the *mhqR^+^* complemented strain and the *mhqR* variants mutated in **(A)** the ligand pocket (F11A, F39A, E43A, H111A and E43A,H111A) and **(B-D)** the predicted DNA-interaction sites (α2/α3: E38A, Q54A), (α3-α4 loop: A63E, S64A, S65A, S66A), (α4: S68A, Y69A, V70A, Q73A) and the β-hairpin (D88A, K89A, R90A, V91A, Y92A). Strains were grown in RPMI to an OD_500_ of 0.5 and harvested before (-) and 30 min after exposure to 100 µM MHQ (+). The arrows point toward the *mhqRED* specific mRNA. The methylene blue stain of the 16S and 23S rRNAs is shown as RNA loading control below the Northern blots. Northern blots were performed from 3 biological replicates and one representative is shown. Using qRT-PCR analysis we determined the relative transcriptional level of *mhqE*, which was normalized to that of untreated WT cells (set to a fold-change of 1). Error bars represent the standard deviation of 3 biological replicates with 3 technical replicates each. Statistical differences of MHQ-induction and derepression of *mhqRED* transcription were determined using two-tailed t-test of two samples of unequal variance, including the comparison co/MHQ, co variants/co *mhqR*^+^ and co variants/co Δ*mhqR*. \**p* ≤ 0.05; \*\**p* ≤ 0.01 and \*\*\**p* ≤ 0.001.

We further investigated the basal transcription and MHQ response of the *mhqRED* operon in *S. aureus* strains expressing the *Sa*MhqR variants with mutations in the predicted DNA-interaction sites *in vivo* to reveal their roles in DNA binding **(Fig. 7B-D)**. The Northern blot results showed that the *Sa*MhqR variants with mutations in predicted DNA binding residues, including the critical α3-α4 allosteric loop (A63E, S65A, S66A), the helices α2 (E38A, Q54A) and α4 (Q73A) and in the ß1/ß2-wing (D88A, K89A, V91A, Y92A) were impaired in DNA binding *in vivo*, as revealed by derepression of transcription of the *mhqRED* operon under control conditions as compared to the *mhqR*^+^ strain **(Fig. 7B-D)**. While most of these non-binding mutants showed no significant MHQ response, the level of derepression under control conditions varied strongly. Quantification using qRT-PCR analysis of all mutants with impaired DNA binding activity revealed that the E38A mutant in helix α2, S65A and S66A in the α3-α4 allosteric loop and the ß1/ß2-wing mutants D88A, K89A, V91A and Y92A showed the full derepression under control conditions as the Δ*mhqR* mutant **(Fig. 7B, D)**. However, we could not validate a significant defect in DNA binding for the variants Q54A, S64A, S68A, Y69A, V70A, Q73A and R90A *in vivo*, indicating that these residues are not essential for DNA binding *in vivo*. Overall, our Northern blot and qRT-PCR results validated the essential role of S65A and S66A of the α3-α4 allosteric loop, and the ß1/ß2-wing residues for the DNA binding activity of *Sa*MhqR, which is consistent with the EMSA results.

Next, we used growth phenotype analyses of the *S. aureus* strains expressing the variants to investigate their MHQ sensitivity **(Fig. S12)**. The phenotype results revealed that the *S. aureus* strains expressing variants in the pocket residues F39A, E43A, H111A, and E43A,H111A were significantly more sensitive in growth towards MHQ compared to the *mhqR*^+^ strain **(Fig. S12)**, which is in agreement with their lack of MHQ response in the transcriptional studies **(Fig. 7A),** supporting their role in MHQ/MBQ-binding as revealed by the crystal structure of the MBQ-bound *Sa*MhqR complex **(Fig. 3C)**. Of the variants predicted to be involved DNA binding, E38A, Q54A, A63E, S66A, D88A, K89A, and Y92A were significantly more resistant to MHQ at few time points along the growth curves, which is consistent with the differential derepression of transcription of the *mhqRED* operon under control conditions in Northern blots, supporting their roles in DNA-binding **(Fig. S12)**.

In conclusion, our DNA-binding studies, transcriptional and growth phenotype analyses confirmed that F11, F39 and H111 are involved in quinone sensing, whereas the S65 and S66 residues of the α3-α4 allosteric loop and the ß1/ß2-wing residues D88A, K89A, V91A, Y92A are essential for DNA binding of *Sa*MhqR, supporting our crystal structure data, models and simulation trajectories upon transition from the DNA-bound to the MHQ-bound *Sa*MhqR protein.

## Discussion

In this work, we determined the crystal structures of the MarR-family regulator *Sa*MhqR and its MBQ-bound complex, providing structural insights into its regulatory mechanism. We further used AlphaFold3 and MD simulation to predict the DNA-bound *Sa*MhqR complex and to elucidate the structural changes and dynamic motions of *Sa*MhqR during transition from the DNA-bound to the quinone-bound conformation. MhqR homologs are widely distributed across Firmicutes and have been previously shown to sense and respond to quinones in *B. subtilis* and *S. aureus* (16, 33). The MhqR homologs of *S. aureus* and *B. subtilis* recognize similar palindromic operator sequences **(Fig. S13)** and control conserved quinone detoxification genes, including dioxygenases and quinone reductases (16, 33). Moreover, sequence alignments of MhqR homologs across Firmicutes, including *Bacillus*, *Staphylococcus*, *Streptococcus* and *Enterococcus* species revealed a similar high conservation with >40% sequence identities **(Fig. S14)** (16). Thus, we have elucidated here the regulatory mechanism of the MhqR family of quinone-responsive MarR-type repressors in Firmicutes.

The structure of the apo *Sa*MhqR repressor resembles that of a typical MarR-type regulator (28–30, 43). Each subunit of the *Sa*MhqR dimer consists of a dimerization domain and a wHTH motif, which is involved in DNA binding. Structural and sequence comparison between the *Sa*MhqR structure and the *Bs*MhqR model revealed a high conservation of the wHTH motifs formed by the helices α2, α3 and α4 and the ß1/ß2-wing in each subunit, supporting the recognition of similar operator sequences (16, 33) **(Fig. 1C, D and Fig. S13)**. Using AlphaFold3, we successfully predicted the structure of the DNA-bound *Sa*MhqR complex to elucidate the interacting residues **(Fig. 2)**. The helix α4 was primarily engaged in base-specific recognition, whereas the α2, α3 helices and the ß1/ß2-wing contribute to non-specific interactions with the sugar-phosphate backbone. Interestingly, the comparison of the apo *Sa*MhqR with DNA-bound *Sa*MhqR revealed a structural rearrangement within the wHTH motifs, especially in the α3-α4 loop region (positions 60-66) upon DNA binding. The apo *Sa*MhqR structure contains an extended unstructured α3-α4 loop with two serine residues S65 and S66, leading to a shorter helix α4 that is unable to interact with the DNA. Upon DNA binding, these serine residues undergo local rearrangements and adapt a helical wound conformation to elongate the helix α4, enabling binding to the major groove of the operator dsDNA sequence. In the predicted DNA-bound form, S65, S66, S68 and Y69 recognize the inverted repeat sequences (TCTCGAA-ATCGAAA) of the *Sa*MhqR operator, while R90 of the ß1/ß2-wing makes additional contacts with minor groove TA bases of the palindromic sequence.

Mutational analyses of predicted DNA-binding residues in the α2, α3 and α4 helices, the α3-α4 loop and the ß1/ß2-wing confirmed that S65 and S66 are essential for DNA binding as revealed by EMSAs *in vitro* and transcriptional analyses *in vivo*. Furthermore, the residues of the ß1/ß2-wing D88, K89, Y90 and V91 were important for DNA binding *in vitro* and *in vivo*, since the mutants showed full derepression of *mhqRED* transcription under control conditions as the Δ*mhqR* deletion mutant. However, lower levels of derepression of *mhqRED* transcription under control conditions and MHQ resistant phenotypes were found in the E38A, Q54A and A63E mutants using Northern blots and growth assays, indicating that these predicted DNA-binding residues are partially required for DNA binding *in vivo*. Our detailed mutational analyses of predicted DNA-binding residues validated several essential DNA-binding residues in the α3-α4 loop and the ß1/ß2-wing, whereas other predicted DNA contacts are less important as compared to those crucial for the structural and conformational change.

Interestingly, this mechanism of transcriptional repression via an allosteric α3-α4 loop region has been recently described for the UrtR family of uric acid-responsive regulators, such as PecS, MftR and HucR (1). Also in PecS, a similar T/S-S-G-T core element within this allosteric loop was identified as essential for major groove binding with high conservation throughout the UrtR family (1). Sequence comparison of MhqR homologs across different Firmicutes confirmed the high conservation of the α3-α4 allosteric loop (L_61_IASSSI_67_) as well as S68 and Y69, indicating a conserved mode of DNA binding of the MhqR family via the helical wound conformation of the allosteric S-S/G-S region **(Fig. S14)**.

Additionally, as in UrtR proteins (1), this allosteric loop region is involved in the regulatory mechanism of *Sa*MhqR by quinones as revealed by the crystal structure of the *Sa*MhqR-MBQ complex, which was successfully resolved in our work **(Fig. 4)**. The MBQ-binding pocket involves residues F11, F39, E43 and H111, which are conserved across other ligand-binding pockets of MarR proteins, as for example in Rv2887 of *M. tuberculosis* (37, 38), ST1710 of *S. tokodaii* (39), AbsC of *S. coelicolor* (40), and HucR of *D. radiodurans* (41, 42) **(Fig. S5)**. However, mutational analysis of F11A, F39A, E43A and H111A variants did not reveal different MHQ and MBQ binding affinities compared to the WT *Sa*MhqR protein based on our ITC experiments *in vitro*. MBQ-binding of single mutants in the pocket residues might be caused by compensatory ligand-binding residues. However, the E43A,H111A double mutant was unable to bind to the DNA in EMSAs and therefore could not be analyzed for MBQ-binding.

In contrast to the *in vitro* EMSA results, transcriptional studies and growth phenotype analyses provided strong evidence that the quinone-pocket residues F11, F39, and H111 are required for sensing of MHQ *in vivo*, since the variants were impaired in the MHQ response and sensitive to MHQ stress. Transcriptional studies showed an even stronger basal repression of the *mhqRED* operon for the H111A and E43A,H111A variants than for the WT protein in the *mhqR*^+^ strain. This might point to a stabilization of the extended α4 for improved DNA binding in the H111A variant. It might be also possible that the non-bound quinone fraction leads to higher induction of the QsrR regulon, compensating for the absence of the MhqR response in the H111A variant. The possible cross-talk of the QsrR and MhqR regulons will be investigated in our future studies. Altogether, the transcriptional data of the variants support our crystal structure of the MhqR-MBQ complex, showing specific contacts of E43 and H111 with the carbonyl group of MBQ, while the phenyl side chains of F11 and F39 possibly interact via π-stacking with the quinone ligand. Interestingly, while the conserved His111 is essential for the binding of MBQ to *Sa*MhqR, its homolog residue His142 is involved in uric acid binding by the UrtR-family HucR repressor (1). This His142 residue of HucR was also shown to form hydrogen bonds with the carbonyl group of uric acid (1), indicating the crucial role of the histidine residue as contact site for carbonyl oxygen in MarR-type proteins, which bind phenolic ligands. Our ligand-bound crystal structure provided also evidence that MhqR most likely binds oxidized benzoquinones, occurring during autoxidation of MHQ, as supported by the fact that the MhqR homolog of *B. subtilis* controls a quinone reductase (AzoR2), which functions in quinone reduction to hydroquinone (33). Additionally, in HucR, residues in the α3-α4 allosteric loop region (T90, M92 and V93) were shown to participate in ligand binding and stabilized the extended coil of the allosteric loop, to prevent the formation of the elongated α4 helix for DNA-binding (1). Similarly, the α3-α4 allosteric loop in *Sa*MhqR also participates in DNA binding by formation of the elongated α4 helix via Ser65 and Ser66. This helical wound conformation of these serine residues is prevented upon MBQ binding, leading to an extended coil of the allosteric loop, which induces a steric clash with the DNA, resulting in dissociation from the operator. However, despite similar DNA binding and regulatory mechanisms involving the allosteric α3-α4 loop region, the ligand pocket residues F11, F39 and E43 of *Sa*MhqR are not conserved in the UrtR family (1), indicating specific interactions of these MarR-type regulators with their specific ligands. Apart from this structural change of the α3-α4 allosteric loop from the helical wound to the unstructured coil conformation during MHQ/MBQ binding, we applied coarse-grained simulations to model the intrinsic dynamics during transition from the DNA-to MHQ-bound states. The simulations revealed a high increase of the intrinsic dynamics of *Sa*MhqR upon MHQ-binding especially in the ß1/ß2-wing, which is required for minor groove DNA binding. Thus, our results revealed that both the structural changes in the allosteric α3-α4 loop and the high intrinsic dynamics of the ß1/ß2-wing contribute to prevent DNA binding of the MHQ-bound conformation.

Altogether, our crystal structure of the quinone-bound *Sa*MhqR complex and the structure-guided model of DNA-bound *Sa*MhqR, which is supported by MD simulations of the intermediate conformations and sampled motions, provides novel insights into the DNA-binding mechanism and regulation of the conserved MhqR family of quinone-sensing MarR-type regulators. This MhqR family has been characterized in *S. aureus* and *B. subtilis* with similar functions in quinone detoxification and antimicrobial resistance. Thus, the quinone-binding site in *Sa*MhqR can be exploited for future drug discovery to combat multi-drug resistance in this important human pathogen *S. aureus*.

## Material & Methods

### Bacterial strains and cultivations

Bacterial strains, plasmids and primers are listed in **Tables S3 and S4**. *Escherichia coli* strains were cultivated in Luria broth (LB) for plasmid construction and protein expression. For stress experiments, *S. aureus* strains were cultivated in RPMI medium supplemented with FeCl_2_ and L-glutamine and treated with 50 µM MHQ at an optical density at 500 nm (OD_500_) of 0.5 as described previously (16). The statistics of the growth curves and the qRT-PCR results were calculated using Student’s unpaired two-tailed t-test.

### Cloning of His-tagged *Sa*MhqR variants in *E. coli*

The *Sa*MhqR alleles F11A was PCR amplified from chromosomal DNA of *S. aureus* COL using primers pET-mhqRF11A-for-NheI and pET-mhqR-rev-BamHI **(Table S4)**. The PCR products were digested with *Nhe*I and *Bam*HI and cloned into pET11b plasmids. The pET11b plasmids expressing the *Sa*MhqR alleles F39A, E43A, E38A, Q54A, A63E, S64A, S65A, S66A, S68A, Y69A, V70A, Q73A, D88A, K89A, V91A, Y92A, H111A and E43A,H111A were constructed using PCR mutagenesis with primers including the alanine substitutions as previously described (26). Primers were used in two first-round PCRs for each *Sa*MhqR variant as listed in **Table S4**. The PCR products of each construct were fused by overlap extension PCR with primers pET-mhqR-for-NheI and pET-mhqR-rev-BamHI to generate the corresponding mutant allele. The PCR products were digested with *Nhe*I and *Bam*HI and cloned into plasmid pET11b, to obtain the pET11b plasmids encoding the variants F39A, E43A, E38A, Q54A, A63E, S64A, S65A, S66A, S68A, Y69A, V70A, Q73A, D88A, K89A, V91A, Y92A, H111A and E43A,H111A. The mutations of the cloned *mhqR* variants were validated by sequencing.

### Protein expression and purification of *Sa*MhqR and variants for EMSAs

For expression and purification of His-tagged *Sa*MhqR WT and *Sa*MhqR variants, *E. coli* BL21(DE3) pLysS was transformed with the plasmids pET11b encoding the *Sa*MhqR variants. Strains were cultivated in 1 Liter LB medium until an OD_600_ of 0.6 followed by addition of 1 mM isopropyl-β-D-thiogalactopyranoside (IPTG) for 3.5 h at 37°C. Recombinant His-tagged proteins were purified using His Trap™ HP Ni-NTA columns and the ÄKTA purifier liquid chromatography system as described (26).

### Protein expression and purification of *Sa*MhqR and variants for protein crystallization and ITC assays

Chemically competent *E. coli* BL21 (DE3) cells (Novagen) were transformed with the pET11b plasmids encoding the C-terminally His-tagged *Sa*MhqR or *Sa*MhqR variant proteins. The strains were grown in 2 liters of auto-inductive Luria-Miller broth (Roth) containing 1% (w/v) α-lactose (Roth) for 20 h at 30 °C and 180 rpm. The cells were subsequently harvested by centrifugation (4,000x*g*, 15 min, 4 °C), resuspended in HEPES buffer (20 mM HEPES, 20 mM MgCl_2_, 200 mM NaCl, 20 mM KCl, 40 mM imidazole, pH 8.0), and disrupted using a microfluidizer (M110-L, Microfluidics). The cell debris was removed by centrifugation (50,000x*g*, 20 min, 4 °C). The supernatant was loaded onto Ni-NTA FF-HisTrap columns (GE Healthcare) for affinity purification. The columns were washed with HEPES buffer (10x column volume) and eluted with HEPES buffer containing 250 mM imidazole. The protein was concentrated with Amicon Ultra-30K centrifugal filters and subjected to size exclusion chromatography (SEC) using a HiLoad 16/600 or HiLoad 26/600 Superdex 200 column equilibrated in 0.2-μm-filtered HEPES buffer without imidazole and a pH of 7.5. The peak fractions were analyzed using a standard SDS-PAGE protocol, pooled, concentrated with Amicon Ultra-30K centrifugal filters, and either used immediately for crystallization experiments, or flash frozen in liquid nitrogen, and stored at -80°C for subsequent ITC experiments. Protein concentrations were determined using a NanoDrop One spectrophotometer (Thermo Fisher Scientific) and respective extinction coefficients.

### Isothermal titration calorimetry (ITC)

The thermodynamic parameters for the binding interaction between *Sa*MhqR and MHQ were assessed using a MicroCal PEAQ-ITC instrument (Malvern Panalytical). Protein concentration was determined using a NanoDrop One spectrophotometer (Thermo Fisher Scientific), while the ligand was dissolved to prepare a 1 M stock solution in water. 20 µM or 50 µM of protein and 500 µM of ligand were prepared in 0.2-μm-filtered SEC buffer (20 mM HEPES, pH 7.5; 200 mM NaCl; 20 mM MgCl_2_, and 20 mM KCl) and loaded into the cell and syringe, respectively. The experiments were conducted at 25 °C with a stirring speed of 750 rpm. Following a pre-injection of 0.4 µL, 12 or 18 injections of 3 µL were made at 150-second intervals. Data analysis was performed using the MicroCal PEAQ-ITC Analysis Software version 1.41 (Malvern Panalytical) with a One Set of Sites model as well as graphically displayed using GraphPad Prism. The ITC assays were performed from two replicates and the resulting thermodynamic values are presented in **Table S2**.

### Structural analysis of apo *Sa*MhqR using X-ray crystallography

Crystallization of *Sa*MhqR WT protein was performed by the hanging-drop method at 20 °C in 1 – 2 µl drops, consisting of protein and precipitation solutions in a 1:1 or 1:2 ratio. Apo-*Sa*MhqR crystallized at a concentration of 2 mM within 1 day in 0.1 M phosphate citrate with 25 – 30% PEG 300 at pH 4.0. Co-crystallization of *Sa*MhqR at a concentration of 2 mM in the presence of 10 mM MHQ was achieved in a crystallization condition containing 1.0 - 1.4 M tri-Sodium citrate and 0.1 M Sodium HEPES pH 7.5. Yellow crystals appeared within 1 day, indicating that the bound ligand had undergone oxidation to MBQ. Before data collection, *Sa*MhqR-MBQ crystals were soaked in crystallization solution supplemented with 20% (v/v) glycerol and flash-frozen in liquid nitrogen. Crystals of apo-*Sa*MhqR were directly flash-frozen in liquid nitrogen. Synchrotron data was collected under cryogenic conditions at the ID23A-1 beamline for the apo-*Sa*MhqR and the ID30A-3 for *Sa*MhqR in complex with MBQ. Both beamlines are operated by the European Synchrotron Radiation Facility (ESRF) in Grenoble, France. X-ray data was processed with XDS and XSCALE (44). Initial phases were determined by molecular replacement using PHASER (45) and a truncated AlphaFold3 model (34) of SaMhqR as search template. Final structures were obtained through iterative model building in COOT (46) and refinement using PHENIX (47). The final structures of *Sa*MhqR were uploaded to the RCSB PDB under accession number 9QDR (apo) and 9SMZ (MBQ-bound). Data were rendered and visualized with UCSF ChimeraX (v.1.10) (48).

### Size exclusion coupled multi-angle light scattering (SEC-MALS)

SEC-MALS was performed using an Äkta PURE system (GE Healthcare) with a Superdex 200 Increase 10/300 column attached to a MALS detector 3609 (Postnova Analytics) and a refractive index detector 3150 (Postnova Analytics). The column was equilibrated with 0.2-μm-filtered HEPES buffer (20 mM HEPES, 20 mM MgCl_2_, 200 mM NaCl, 20 mM KCl). The molecular weight was calculated by combining the refraction index and MALS values using Zimmermann fitting.

### Modeling of proteins and protein complexes using AlphaFold3

Protein and protein–DNA complex structures were predicted using the AlphaFold3 server provided by Google DeepMind (alphafoldserver.com) (34). Compared to its predecessor AlphaFold3 is using a substantially revised diffusion-based architecture capable of predicting joint structures of complexes comprising proteins, nucleic acids, small molecules, ions, and modified residues. We submitted amino acid and DNA sequences in FASTA format through the web interface, and applied default parameters for prediction. Generated models were downloaded in PDB format and evaluated based on the reported confidence metrics using UCSF ChimeraX (48). Namely, pLDDT (predicted Local Distance Difference Test) indicating the per-residue confidence score, ipTM (interface predicted TM-score) for the confidence of inter-chain interfaces in complexes and pTM (predicted TM-score) as the overall model topology confidence.

### Molecular dynamic (MD) simulations

MD simulations were carried out using the *Sa*MhqR structures apo *Sa*MhqR dimer, DNA-bound and MHQ-bound *Sa*MhqR dimers, and apo *Sa*MhqR monomer (as control). The *Sa*MhqR monomer was modelled using AlphaFold3 (34). All structures were visually inspected, and prepared using UCSF ChimeraX (48). The preparation consisted of molecular-mechanical energy minimization (up to 1,000 steps using steepest-descent algorithm). Energy-minimized structures were checked and imported into GROMACS (49) for atomistic (all-atom) MD simulations.

### Atomistic MD simulations

All simulations used GROMACS (49) with AMBER99SB-ILDN (50) force field. Each simulated structure was immersed in a cubic TIP3P water box, set to be 1 nm away from the edge of the protein. 3D-PBC were applied. Na^+^ and Cl^−^ ions were added to maintain the charge neutrality of simulated units. Prior MD simulations, each solvated system underwent energy minimization via the steepest descent method for 1,000 cycles and the conjugate gradient method for further refinement, with the energy step size of 0.001 nm and a maximum of 50,000 steps. The minimization was concluded when the maximal force descended below 1000 kJ/mol/nm. Long-range electrostatic interactions were addressed using the Particle-Mesh Ewald (PME) method (51) while a cut-off of 1.0 nm was applied for short-range electrostatic and van der Waals interactions. Following energy minimization, all systems were subjected to a 500 ps NVT equilibration with a step size of 2 fs. The protein and non-protein groups were gradually heated to the target temperature (300 K) under the influence of the V-rescale thermostat with a time constant of 0.1 ps. LINCS (Linear Constraint Solver) position restraints were applied to the bond lengths and angles of the backbone atoms. The non-bonded short-range interactions were treated with the Verlet cut-off scheme, setting the cut-off distance to 1.0 nm. Long-range electrostatics were once again addressed with PME. Subsequently, NPT equilibration was performed, where temperature was maintained at the target with the continued utilization of the temperature coupler, followed by the initiation of Parrinello-Rahman pressure coupling (52) for 500 ps of pressure equilibration, with the target pressure established at 1 bar. Post-equilibration, the systems were subjected to a single-replica 100 ns simulations to obtain trajectories for analysis, discarding the initial 10 ns of data during the final evaluation. Trajectory analyses were conducted using GROMACS tools. The overall stability was evaluated by root mean square deviation (RMSD), and local flexibility was assessed via per-residue root mean square fluctuation (RMSF). Principal component analysis (PCA) was employed to explore the key motion modes. Secondary structure changes were quantified by DSSP, visualized using a custom in-house script. The protein overall structural stability was analyzed through the radius of gyration (Rg). Examination of hydrogen-bonding network was carried out utilizing UCSF ChimeraX (48). Molecular interactions and binding affinities were estimated using SeeSAR (https://www.biosolveit.de/SeeSAR).

### Coarse-grained simulations

To assess conformational changes induced by MHQ binding to *Sa*MhqR and to predict protein intermediate conformational states, “invisible” to experimental techniques, we used in-house version of Elastic network-driven Brownian Dynamics IMportance Sampling (eBDIMS) (53) code. The workflow involved coarse-grained simulations on two distinct conformers of the *Sa*MhqR dimer: DNA-bound and MHQ-bound. This process enabled the generation of a transition pathway between these two conformers. Next, Normal Mode Analysis (NMA) was applied to obtain the modes and frequencies of protein motions. NMA selected the lowest-frequency motions as the transition pathway. Subsequently, the code utilizing PD2 (54) and Scwrl4 (55) performed reverse mapping on the whole coarse-grained trajectory, resulting in the generation of protein structures at the full atomic level. 20 intermediates were generated for each conformational transition of *Sa*MhqR (DNA-bound to MHQ-bound and MHQ-bound to DNA-bound, respectively).

### Construction of the *S. aureus* COL strains expressing the *Sa*MhqR variants

The coding sequences of the *mhqR* variants (F39A, E43A, E38A, Q54A, A63E, S64A, S65A, S66A, S68A, Y69A, V70A, Q73A, D88A, K89A, V91A, Y92A, H111A and E43A,H111A) were amplified from the corresponding pET11b plasmids using primers mhqR-pRB-for-BamHI and mhqR-pRB-rev-kpnI **(Table S4)**. The purified PCR products were digested with *Bam*HI and *Kpn*I and cloned into plasmid pRB473, resulting in pRB473 plasmids expressing the *Sa*MhqR variants F39A, E43A, E38A, Q54A, A63E, S64A, S65A, S66A, S68A, Y69A, V70A, Q73A, D88A, K89A, V91A, Y92A, H111A and E43A,H111A under control of the xylose-inducible promoter **(Table S3)**. The plasmids were introduced into the *S. aureus* Δ*mhqR* mutant via phage transduction as described (26).

### RNA isolation, Northern blot and quantitative real-time PCR (qRT-PCR) analysis

The *S. aureus* COL strains expressing the *Sa*MhqR variants were cultivated in RPMI with 1% (w/v) xylose and 10 µg/ml chloramphenicol and treated with 100 µM MHQ at an OD_500_ of 0.5 for 30 min as previously described (16). Northern blot hybridizations were conducted with digoxigenin-labeled *mhqE*- and *mhqR*-specific antisense RNA probes, which were synthesized by *in vitro* transcription using T7 RNA polymerase and the primer pairs NB-mhqE-for/rev and NB-mhqR-for/rev **(Table S4)**, as described previously (16, 26). The Northern blot analyses were performed from RNA samples of three biological replicates.

For qRT-PCR analysis, the purified total RNA was reverse-transcribed into cDNA using the RevertAid Reverse Transcriptase (Thermo Fisher Scientific) according to the recommendations of the manufacturer. qRT-PCR analyses were performed using Takyon™ ROX SYBR 2X MasterMix dTTP blue (Eurogentec: UF-RSMT-B0701) and an iQ5 cycler detection system (Bio-Rad; 170-9780). Each reaction had a final volume of 10 μl, including 5 μl SYBR 2X MasterMix, 4 μl of the 10-fold diluted cDNA, 0.2 μl of each primer (10 μM), and water. Reaction conditions included one cycle of 95°C for 3 min, followed by 40 cycles of 95°C for 3 s and 58°C for 30 s. The Ct values from the input samples were obtained in three technical replicates and the fold change in the target gene expression was calculated as previously described (56, 57). The qRT-PCR analyses were performed from RNA samples of three biological replicates with each three technical replicates.

### Electrophoretic mobility shift assays (EMSAs) of purified *Sa*MhqR and *Sa*MhqR variants

For DNA binding assays *in vitro*, EMSAs were performed with the DNA promoter probe as described previously (16, 26). The DNA-binding reactions were performed with 15 ng of the *mhqRED* promoter probe (56), incubated with the purified His-tagged *Sa*MhqR WT or *Sa*MhqR variant proteins for 60 min according to the EMSA protocol as described before (16). To study the effect of MHQ on DNA-binding activity of *Sa*MhqR proteins *in vitro*, the purified proteins were diluted to 0.6 μM in 10 mM Tris-HCl (pH 7.5), treated with 0.5–100 mM MHQ for 30 min at room temperature and incubated with the DNA fragment for 60 min as described above. The EMSA experiments were performed in three replicates.

## ACKNOWLEDGEMENTS

This work was supported by the Deutsche Forschungsgemeinschaft (DFG, Germany) through grants AN746/8-1 and TR84 (project B06) to H.A. P.W. acknowledges support from the DFG Research Training Group GRK 2573 “The inflammatory tumor secretome – from understanding to novel therapies.” (project no. 416910386). F.B. and G.B. acknowledge support from the DFG Research Training Group 2937 “Nucleotide Metabolism in Microbes (MiNu)” (project no. 505997786). We further thank Maurizio Iannuzzi and Beate Koksch (FU Berlin) for assistance with CD measurements. All authors thank the European Synchrotron Radiation Facility (ESRF) in Grenoble, France for excellent support.

## AUTHOR DISCLOSURE STATEMENT

No competing financial interests exist.

## DAT A AV AILABILITY STATEMENT

The authors confirm that the data supporting the findings of this study are available within the article and its supplementary materials. The crystal structure data of the *S. aureus Sa*MhqR proteins are available in the PDB database under accession numbers 9QDR (apo *Sa*MhqR) and 9SMZ (MBQ-bound *Sa*MhqR).

## Notes

### Competing Interest Statement

The authors have declared no competing interest.

