## Supplementary material for "Structural basis of quinone-sensing by the MarR-type repressor MhqR in *Staphylococcus aureus*": Fig.S1-S14; Tables S1-S4

### Figure S1

A

**Hydrogen bonds**

| A | Distance (Å) | B |
|---|---|---|
| ASN 9 | 2.69 | GLN 131 |
| LEU 122 | 2.78 | ARG 3 |
| ARG 3 | 2.79 | LEU 122 |
| ARG 3 | 2.84 | LEU 122 |
| ALA 118 | 2.96 | GLN 140 |
| GLN 131 | 2.97 | ASN 9 |
| LEU 122 | 2.98 | ARG 3 |
| ARG 16 | 2.99 | ARG 59 |
| GLN 131 | 3.00 | GLN 6 |
| GLN 6 | 3.01 | THR 128 |
| GLN 140 | 3.03 | ALA 118 |
| GLN 6 | 3.06 | GLN 131 |
| THR 4 | 3.06 | THR 116 |
| THR 20 | 3.08 | SER 138 |
| LYS 124 | 3.09 | ASP 2 |
| PHE 122 | 3.16 | ARG 3 |
| THR 128 | 3.18 | GLN 6 |
| ARG 3 | 3.28 | PHE 119 |
| LYS 136 | 3.34 | GLU 126 |
| GLU 126 | 3.45 | LYS 136 |
| SER 138 | 3.50 | THR 20 |
| ARG 31 * | 3.63 | HIS 146 * |
| GLN 6 | 3.66 | LYS 124 |
| LYS 124 | 3.73 | GLN 6 |
| GLU 126 | 3.76 | LYS 136 |

**Salt bridges**

| A | Distance (Å) | B |
|---|---|---|
| LYS 124 | 3.09 | ASP 2 |
| LYS 136 | 3.34 | GLU 126 |
| GLU 126 | 3.45 | LYS 136 |
| GLU 126 | 3.76 | LYS 136 |
| LYS 136 | 3.80 | GLU 126 |
| ARG 3 | 3.95 | ASP 120 |

| solvent-accessible area (Å²) | A | B |
|---|---|---|
| Interface | 2890.8 | 2895.1 |
| percent of total | 25.60% | 27.70% |
| **solvation energy, kcal/mol** | | |
| isolated structure | -111.5 | -112.5 |
| gain on complex formation | -20.2 | -21.4 |
| average gain | -10.3 | -10.6 |
| P-value | 0.063 | 0.037 |

B

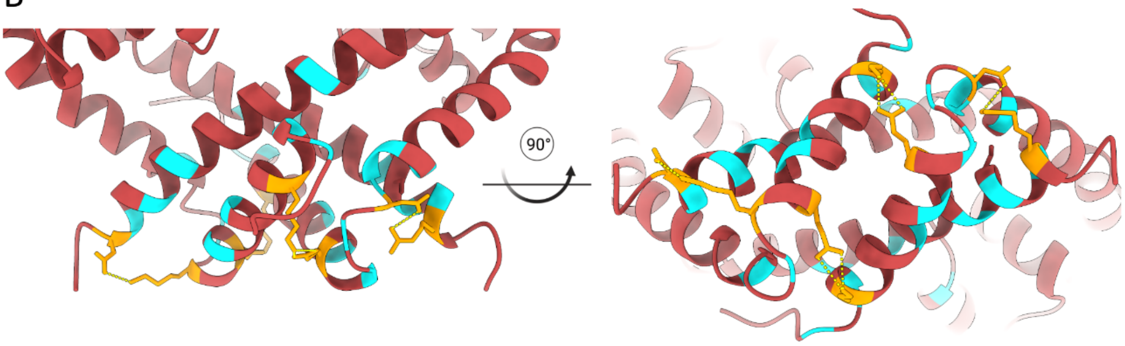


**Figure S1. Description of the *Sa*MhqR dimer interface. A)** Hydrogen bonds (cyan) and salt bridges (yellow) formed by amino acid pairs are listed and sorted based on their distance from each other. Note, His146 (black asterisk) is not part of the natural dimer interface since it is an artifact of the C-terminal hexahistidine tag. The buried interface area (Å²) and its corresponding percentage based on the total surface area are listed together with solvation energies in kcal/mol. **B)** The dimer interface is illustrated with the crystal structure of *Sa*MhqR (red) and hydrogen bonds forming amino acids in cyan and salt bridge forming residues as sticks in yellow. The dimer interface analysis was performed using the PDBe PISA tool (58) developed by the Protein Data Bank in Europe.

#### Figure S2

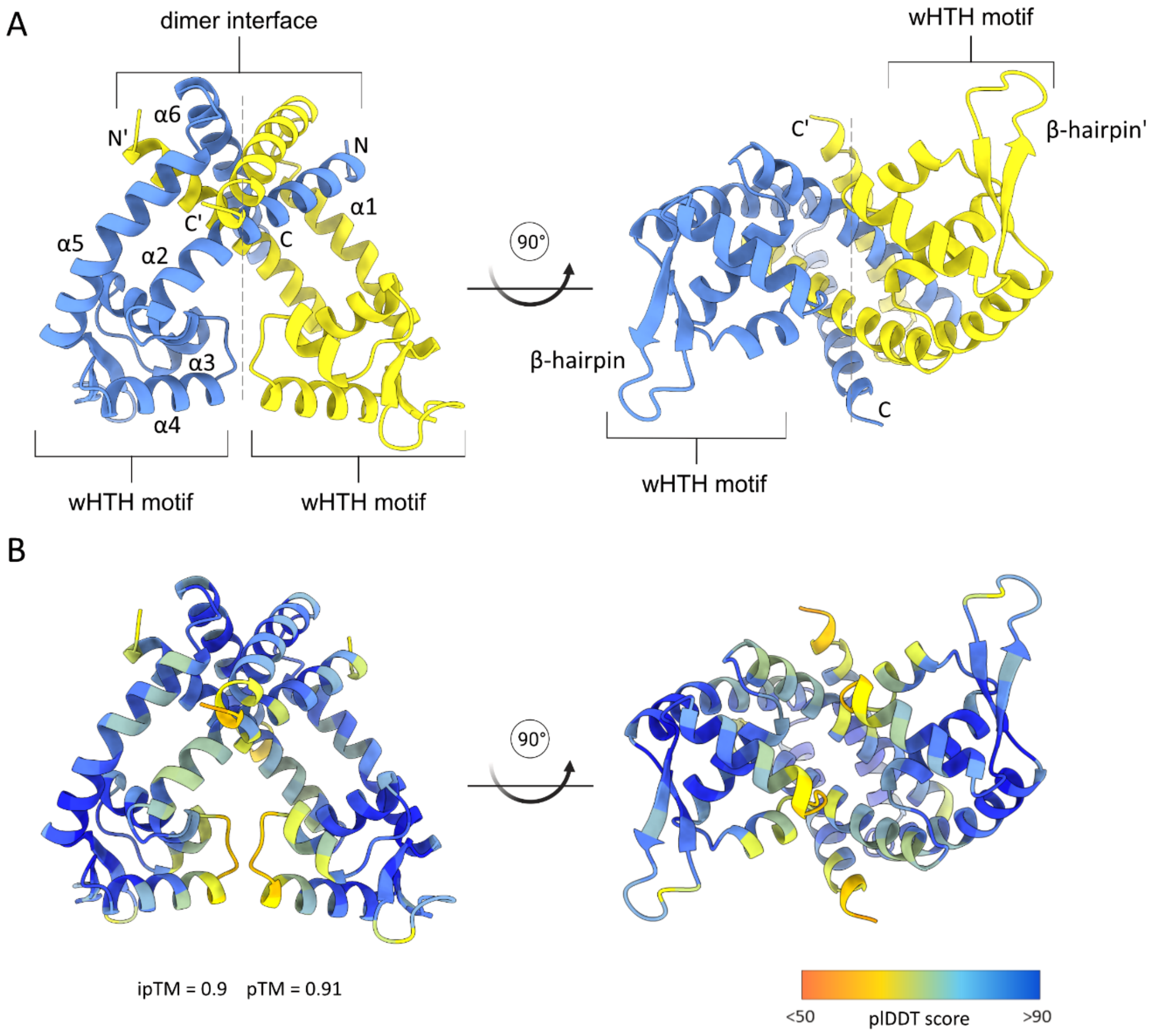


**Figure S2. Prediction of the protein structure of *Bs*MhqR.** The protein structure of *Bs*MhqR was predicted by AlphaFold3 (34) and is shown as side and top views. **(A)** The two monomers of the *Bs*MhqR dimer are colored in yellow and blue. **(B)** The color key of the *Bs*MhqR dimer is based on the AlphaFold palette indicating high confidence (pLDDT score) in blue and low in red.

#### Figure S3

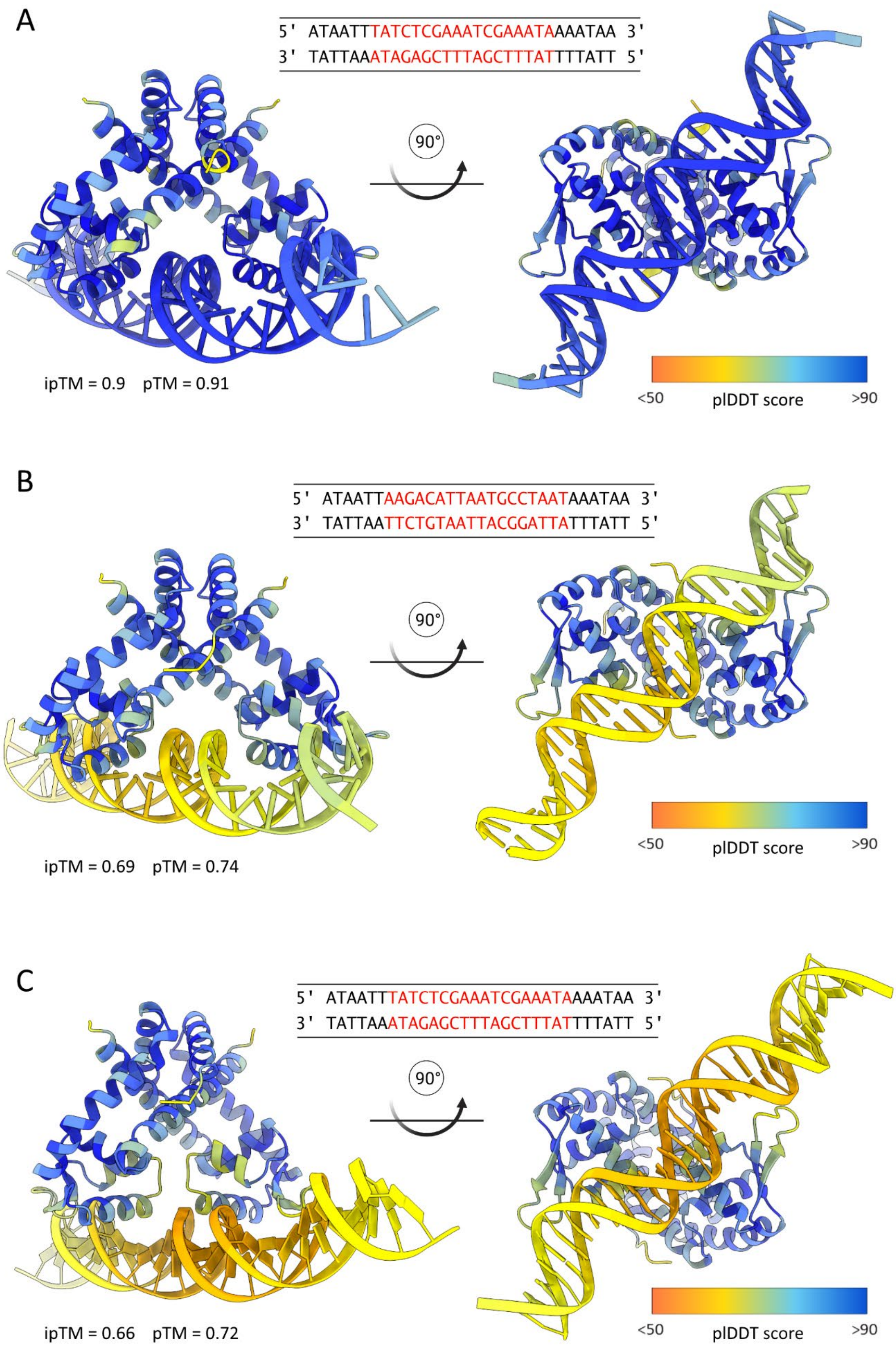
A
5' ATAATTTATCTCGAAATCGAAATAAAATAA 3'
3' TATTAAATAGAGCTTTAGCTTTATTTTATT 5'
90°
ipTM = 0.9 pTM = 0.91
<50
plDDT score
>90
B
5' ATAATTAAGACATTAATGCCTAATAAATAA 3'
3' TATTAATTCTGTAATTACGGATTATTTATT 5'
90°
ipTM = 0.69 pTM = 0.74
<50
plDDT score
>90
C
5' ATAATTTATCTCGAAATCGAAATAAAATAA 3'
3' TATTAAATAGAGCTTTAGCTTTATTTTATT 5'
90°
ipTM = 0.66 pTM = 0.72
<50
plDDT score
>90

**Figure S3. Predicted *Sa*MhqR-DNA complexes. A)** The protein structure of WT *Sa*MhqR bound to its cognate palindromic operator DNA as predicted by AlphaFold3 (34). **B)** The predicted protein structure of *Sa*MhqR bound to a scrambled (non-cognate) operator DNA with the same G-C content. **C)** The predicted protein structure of the *Sa*MhqR S65A, S66A variant bound to its cognate operator DNA. The DNA operator sequence is indicated in red on top of each figure panel. The color key is based on the AlphaFold3 palette indicating high confidence (pLDDT score) in blue and low in red.

#### Figure S4

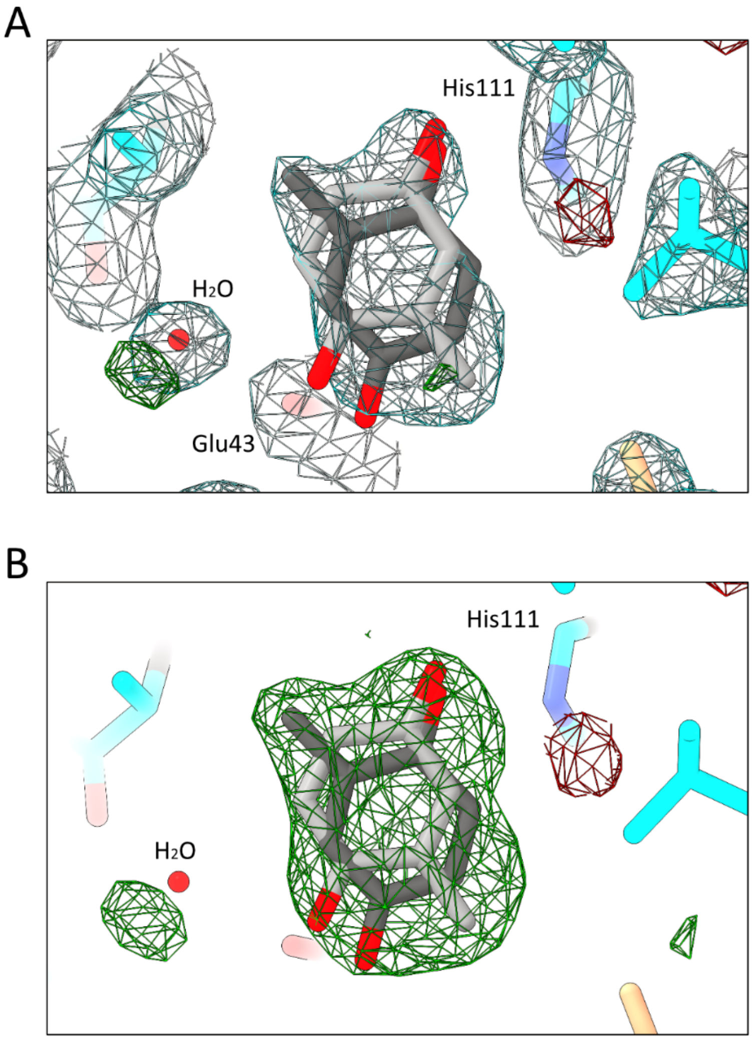


**Figure S4. Electron density maps of *Sa*MhqR in complex with MBQ. A)** The ligand MBQ in two orientations ($q_{1,\text{ light grey}}$ = 0.5; $q_{2,\text{ dark grey}}$ = 0.3) is depicted in the binding pocket with the corresponding 2Fo-Fc map contoured at 1 σ (grey) and the Fo-Fc difference map contoured at 3 σ (green and red). **B)** An Fo-Fc polder omit map, excluding bulk solvent around the omitted region and contoured at 3 σ (green), confirms the presence of MBQ.

#### Figure S5

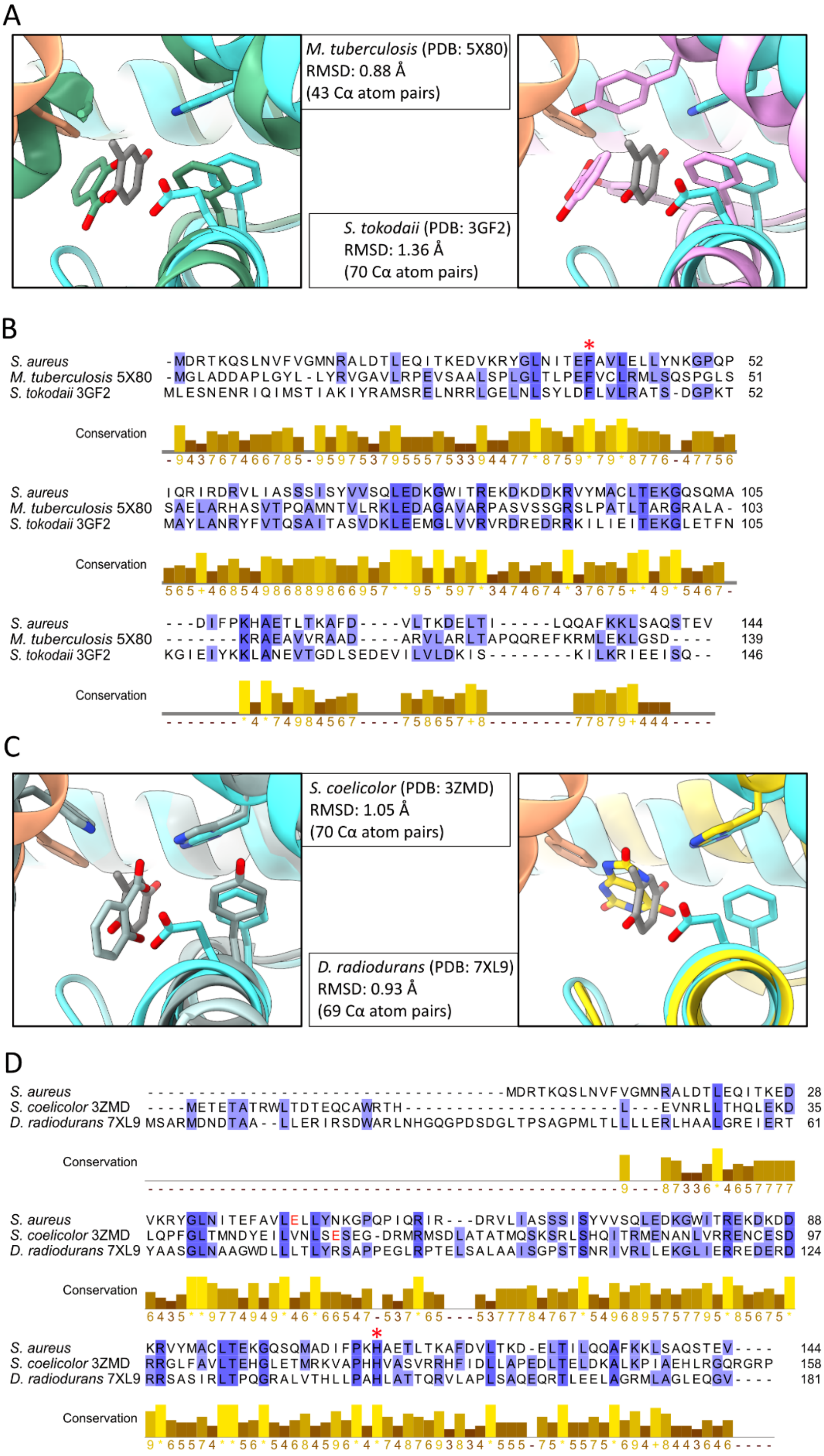

A
M. tuberculosis (PDB: 5X80)
RMSD: 0.88 Å
(43 Cα atom pairs)
S. tokodaii (PDB: 3GF2)
RMSD: 1.36 Å
(70 Cα atom pairs)
B
S. aureus -MDRTKQSLNVFVGMNRALDTLEQITKEDVKRYGLNITEFAVLELLYNKGPQP 52
M. tuberculosis 5X80 -MGLADDAPLGYL-LYRVGAVLRPEVSAALSPLGLTLPEFVCLRMLSQSPGLS 51
S. tokodaii 3GF2 MLESNENRIQIMSTIAKIYRAMSRELNRRLGELNLSYLDFLVLRATS-DGPKT 52
Conservation
S. aureus IQRIRDRVLIASSSISYVVSQLEDKGWITREKDKDDKRVYMACLTEKGQSQMA 105
M. tuberculosis 5X80 SAELARHASVTPQAMNTVLRKLEDAGAVARPASVSSGRSLPATLTARGRALA- 103
S. tokodaii 3GF2 MAYLANRYFVTQSAITASVDKLEEMGLVVRVRDREDRRKILIEITEKGLETFN 105
Conservation
S. aureus ---DIFPKHAETLTKAFD----VLTKDELTI-----LQQAFKKLSAQSTEV 144
M. tuberculosis 5X80 -------KRAEAVVRAAD----ARVLARLTAPQQREFKRMLEKLGSD---- 139
S. tokodaii 3GF2 KGIEIYKKLANEVTGDLSEDEVILVLDKIS--------KILKRIEEISQ-- 146
Conservation
C
S. coelicolor (PDB: 3ZMD)
RMSD: 1.05 Å
(70 Cα atom pairs)
D. radiodurans (PDB: 7XL9)
RMSD: 0.93 Å
(69 Cα atom pairs)
D
S. aureus 28
S. coelicolor 3ZMD 35
D. radiodurans 7XL9 61
Conservation
S. aureus 88
S. coelicolor 3ZMD 97
D. radiodurans 7XL9 124
Conservation
S. aureus 144
S. coelicolor 3ZMD 158
D. radiodurans 7XL9 181
Conservation

**Figure S5. The quinone-binding pocket of *Sa*MhqR is conserved in other MarR-family proteins. A)** The MBQ-binding pocket of *Sa*MhqR (cyan) is superposed with the ligand-binding sites of *Mycobacterium tuberculosis* Rv2887 bound to salicylic acid (PDB:5X80, green) and *Sulfurisphaera tokodaii* ST1710 also bound to salicylic acid (PDB: 3GF2, pink). The root mean square deviation (RMSD) is used to describe the structural differences between the superposed α-carbon positions (Cα) and the Z-score describes the statistical significance of this pairwise comparison. **B)** Sequence alignments highlight the conserved Phe39 (red asterisk) in *Sa*MhqR, Rv2887 and ST1710 on a structural and sequence level. **C)** The MBQ-binding pocket of *Sa*MhqR is superposed with the ligand binding sites of *Streptomyces coelicolor* AbsC bound to salicylic acid (PDB:3ZMD, silver grey) and of *Deinococcus radiodurans* HucR bound to urate (PDB: 7XL9, yellow). **D)** Sequence alignments highlight the conserved Glu43 (red asterisk) in *Sa*MhqR and AbsC on a structural level and His111 (red asterisk) across *Sa*MhqR, AbsC and HucR on a structural and sequence level.

### Figure S6

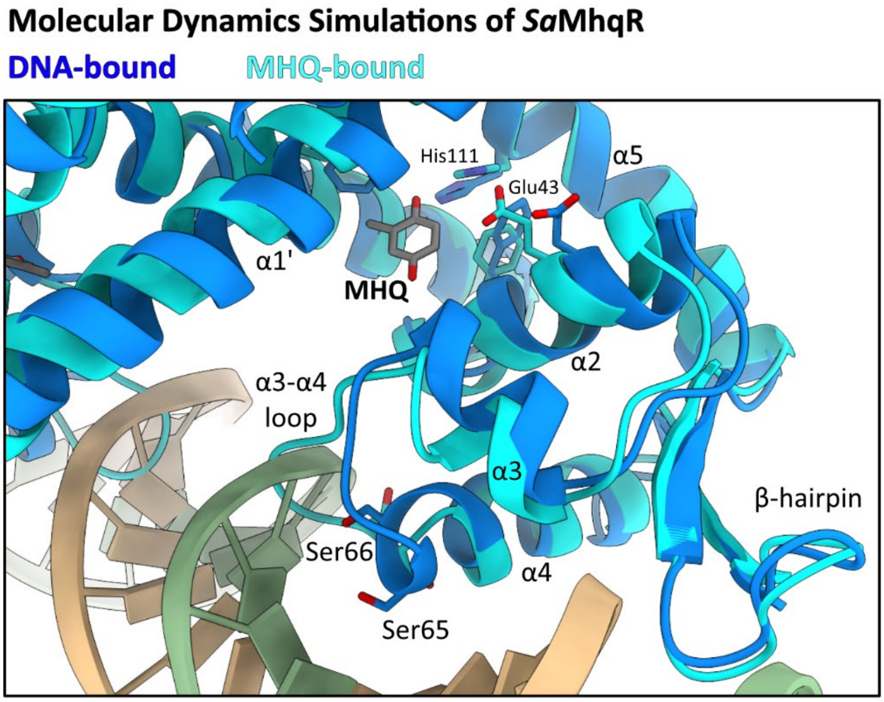


**Figure S6. Conformational changes of the *Sa*MhqR-DNA complex upon MHQ binding after MD simulations.** Superposing the DNA-*Sa*MhqR complex (blue) and the MBQ-bound *Sa*MhqR (cyan), obtained as products of the cluster analyses after MD simulations. The region spanning residues 58-68 of the α3-α4 allosteric loop is shown, highlighting Ser65 and Ser66 that elongate the length α4 and shorten the α3-α4 loop to facilitate DNA binding. Conformational changes induced by MBQ binding causes clash of the allosteric α3-α4 loop with the DNA.

#### Figure S7

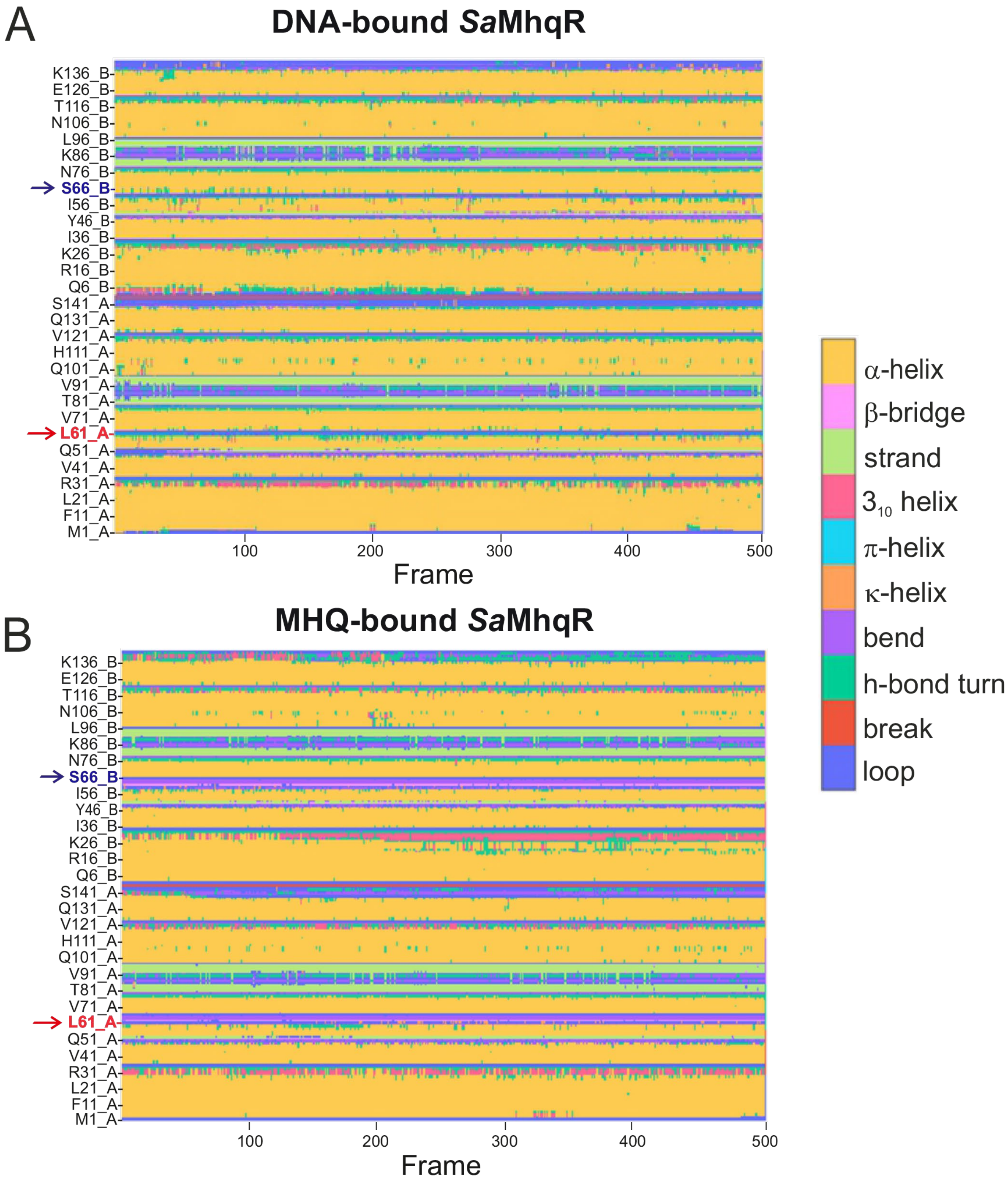


**Figure S7. Secondary structure analysis over MD simulation time.** Change of the secondary structure is shown for every amino acid residue (Y-axis) of **(A)** DNA-bound and **(B)** MHQ-bound *Sa*MhqR complexes, as the MD simulation progresses. The 500 frames (X-axis) were collected during 100 ns of unrestrained MD simulation. Regions which are different in the DNA-bound and MHQ-bound *Sa*MhqR include the α3-α4 loop from Leu61 to Ser66. Ser66 in chain B is marked in blue, and Leu61 in chain A is marked in red.

#### Figure S8

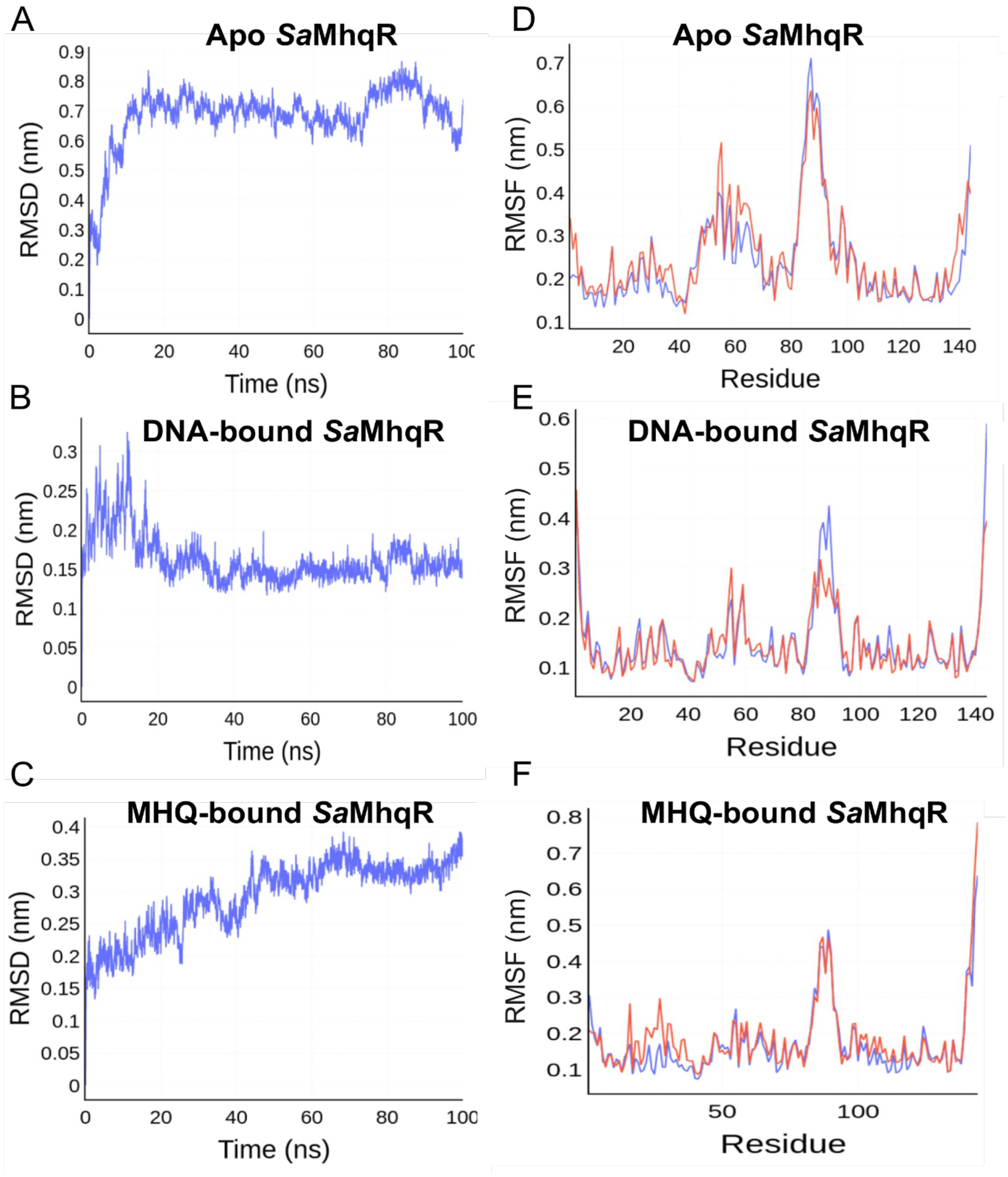


**Figure S8. Root-mean-square deviation (RMSD) and root-mean-square fluctuations (RMSF) of *Sa*MhqR**. RMSD and RMSF are shown for Apo *Sa*MhqR **(A, D)**, DNA-bound **(B, E)**, and MHQ-bound **(C, F)** *Sa*MhqR complexes. RMSD values (Y-axis) are calculated for every non-hydrogen atom in the protein, and displayed as a function of the simulation time (X-axis). RMSF values (Y-axis) are calculated for every residue in the protein (X-axis). The RMSD plots are shown for chain A (blue) and the RMSF plots are shown as overlay of chain A (blue) and chain B (red).

### Figure S9

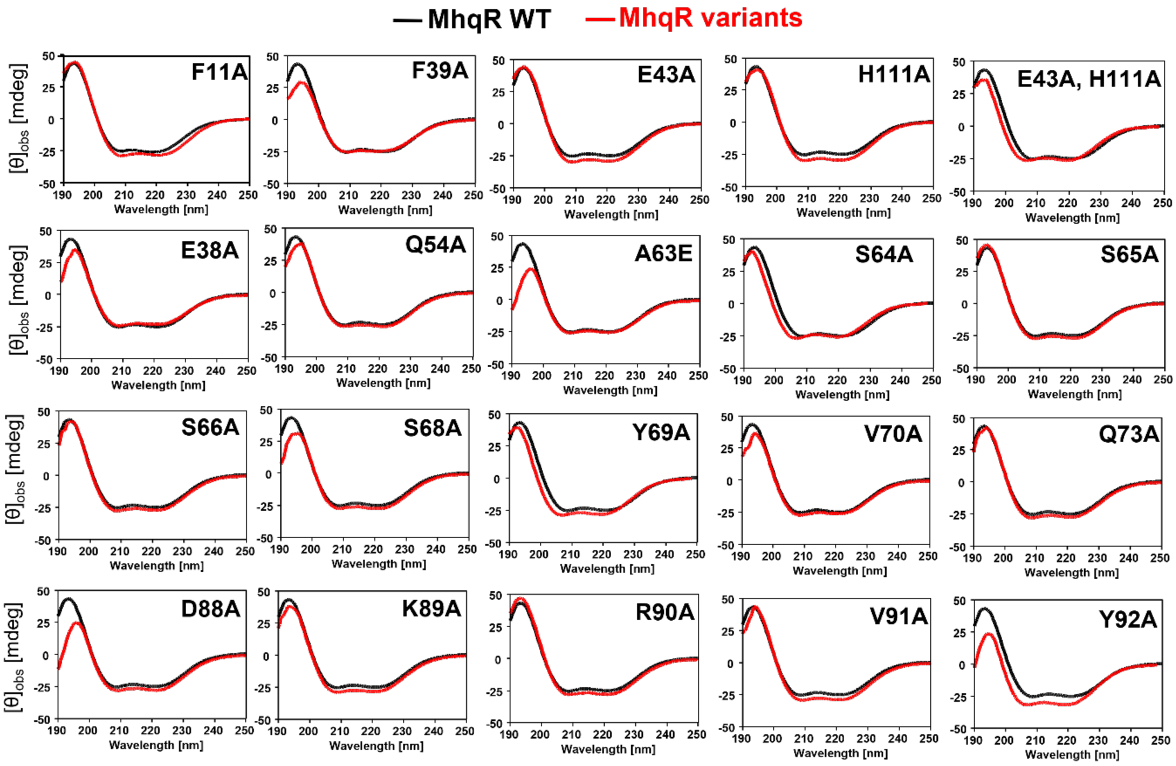


**Figure S9. Circular dichroism (CD) spectra of purified *Sa*MhqR variants.** The CD spectra of the *Sa*MhqR variants (red) and the SaMhqR WT (black) were recorded in the range of 190–250 nm to evaluate changes in secondary structures. CD-spectra were measured using a Jasco J-810 spectropolarimeter at 10 µM protein in 20 mM potassium phosphate buffer, pH 7.5 with 1 mM DTT. $\theta_{obs}$ is the measured ellipticity in degree, as indicated by [mdeg]. The CD measurements with performed from three biological replicates and average values are shown.

### Figure S10

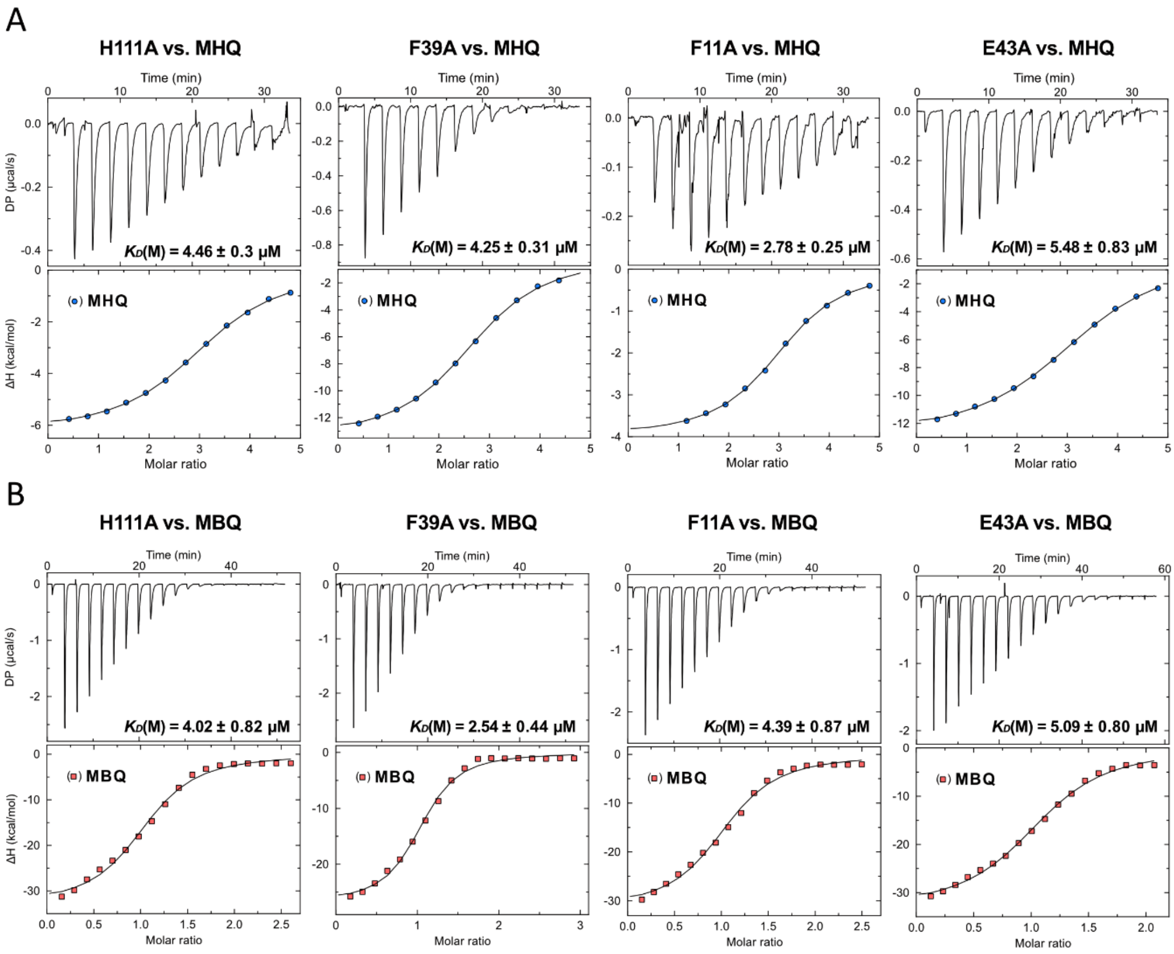


**Figure S10. Isothermal titration calorimetry (ITC) curves of *Sa*MhqR variants.** Fitted ITC curves are determined by ITC experiments with purified *Sa*MhqR variants with mutations in the MBQ-binding pocket (F11A, F39A, E43A, H111A) against **(A)** MHQ and **(B)** MBQ. The ITC experiments were performed in two replicates and one representative replicate is shown for each *Sa*MhqR variant.

#### Figure S11

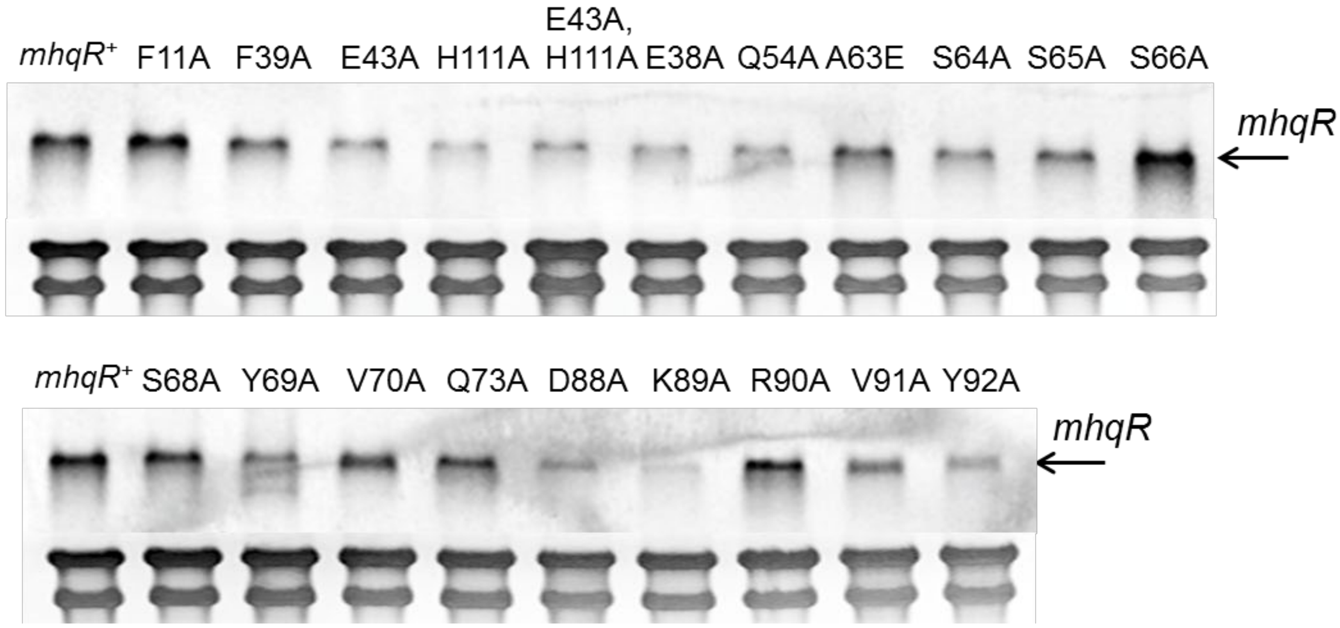


**Figure S11. Northern blot analysis of *mhqR* transcription in *S. aureus* strains expressing *Sa*MhqR variants.** Northern blot analysis of *mhqR* transcription was performed from RNA isolated from *S. aureus* strains expressing the expressing *mhqR*$^+$ or the *mhqR* variants mutated in the ligand pocket (F11A, F39A, E43A, H111A and E43A,H111A) and the predicted DNA-interaction sites (α2/α3: E38, Q54A), (α3-α4 loop: A63E, S64A, S65A, S66A), (α4: S68A, Y69A, V70A, Q73A) and the ß-hairpin (D88A, K89A, V91A, Y92A). Strains were grown in LB containing 1% xylose and harvested at an $OD_{500}$ of 2.0 for RNA isolation. The arrows point toward the *mhqR* specific mRNA, expressed in the variants from plasmid pRB473. The methylene blue image below the Northern blot image denotes the bands of the 16S and 23S rRNA as RNA loading control. The experiments were performed in two biological replicates and one representative is shown.

#### Figure S12

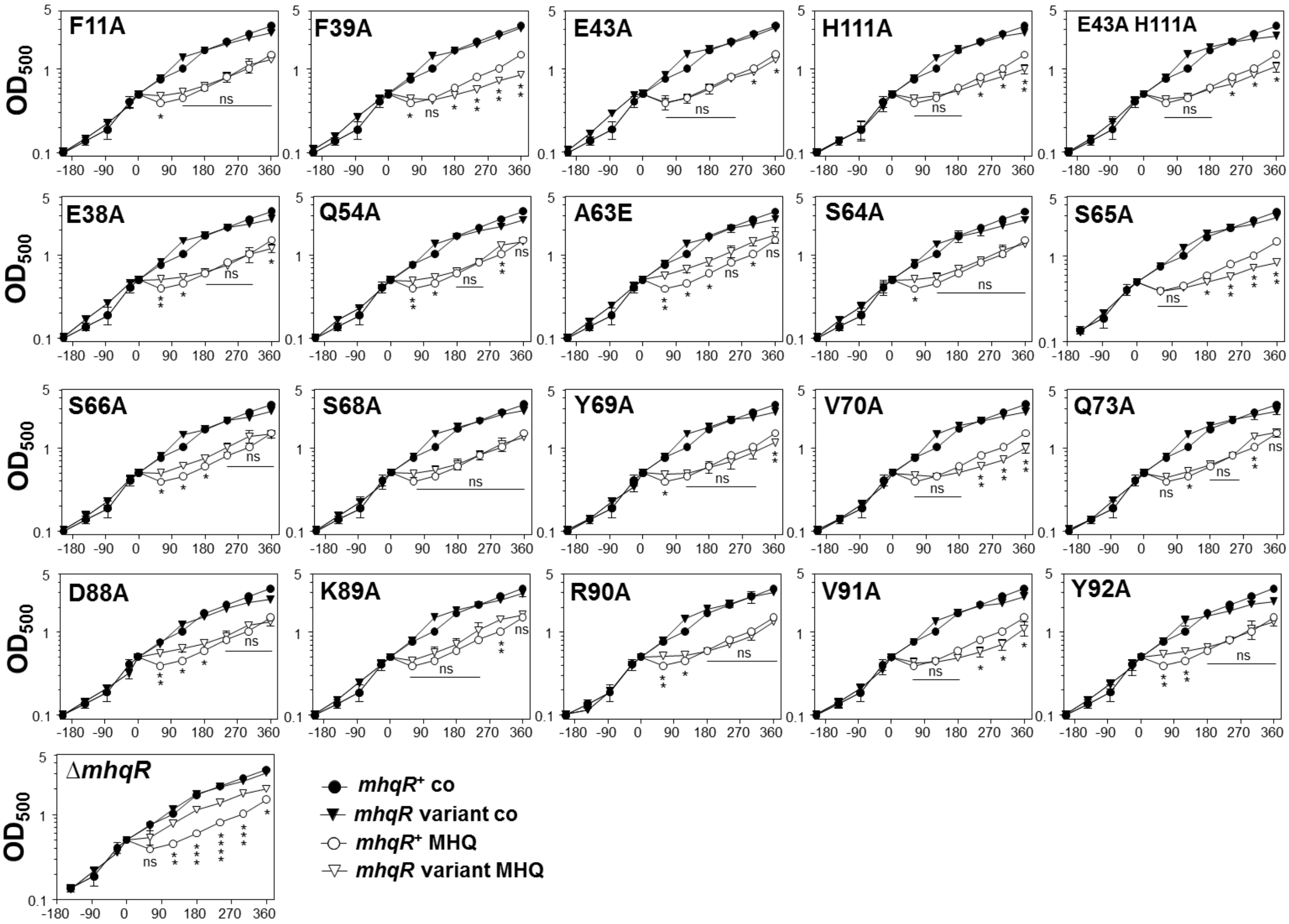


**Figure S12. Growth phenotypes of *S. aureus* strains expressing *Sa*MhqR variants after exposure to MHQ stress.** For growth phenotypes, we compared the MHQ sensitivities of *S. aureus* COL WT, the Δ*mhqR* mutant and complemented strains, expressing *mhqR*$^+$ or the *mhqR* variants mutated in the ligand pocket (F11A, F39A, E43A, H111A and E43A,H111A) and the predicted DNA-interaction sites (α2/α3: E38, Q54A), (α3-α4 loop: A63E, S64A, S65A, S66A), (α4: S68A, Y69A, V70A, Q73A) and the ß-hairpin (D88A, K89A, V91A, Y92A). Strains were grown in RPMI containing 1% xylose until an $OD_{500}$ of 0.5 and treated with 50 µM MHQ to monitor the growth profiles. Mean values were calculated from 3 biological replicates and error bars represent the SD. The statistics of growth differences between the MHQ-treated variants and the Δ*mhqR* mutant *versus* the MHQ-treated *mhqR*$^+$ strain were calculated using a Student's unpaired two-tailed *t*-test for two samples with unequal variance by the GraphPad Prism software. Symbols are: $^{ns}p > 0.05$, $^{*}p \leq 0.05$, $^{**}p \leq 0.01$ and $^{***}p \leq 0.001$.

#### Figure S13

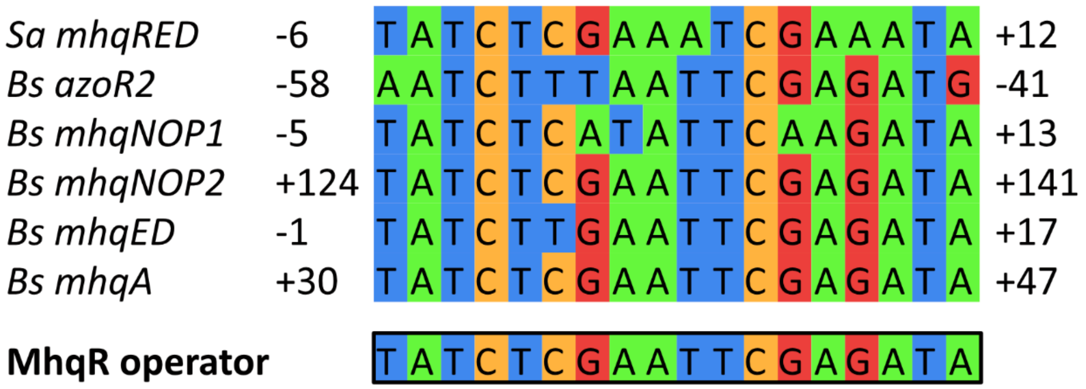


**Figure S13. Alignment of the palindromic MhqR operator sequences upstream of *B. subtilis* and *S. aureus* MhqR regulon members.** The MhqR operator sequence upstream of the *S. aureus mhqRED* operon was aligned with the operators of the *B. subtilis* MhqR regulon members, including the *azoR2, mhqNOP, mhqED* and *mhqA* upstream sequences. The MhqR consensus sequence is indicated at the bottom.

### Figure S14

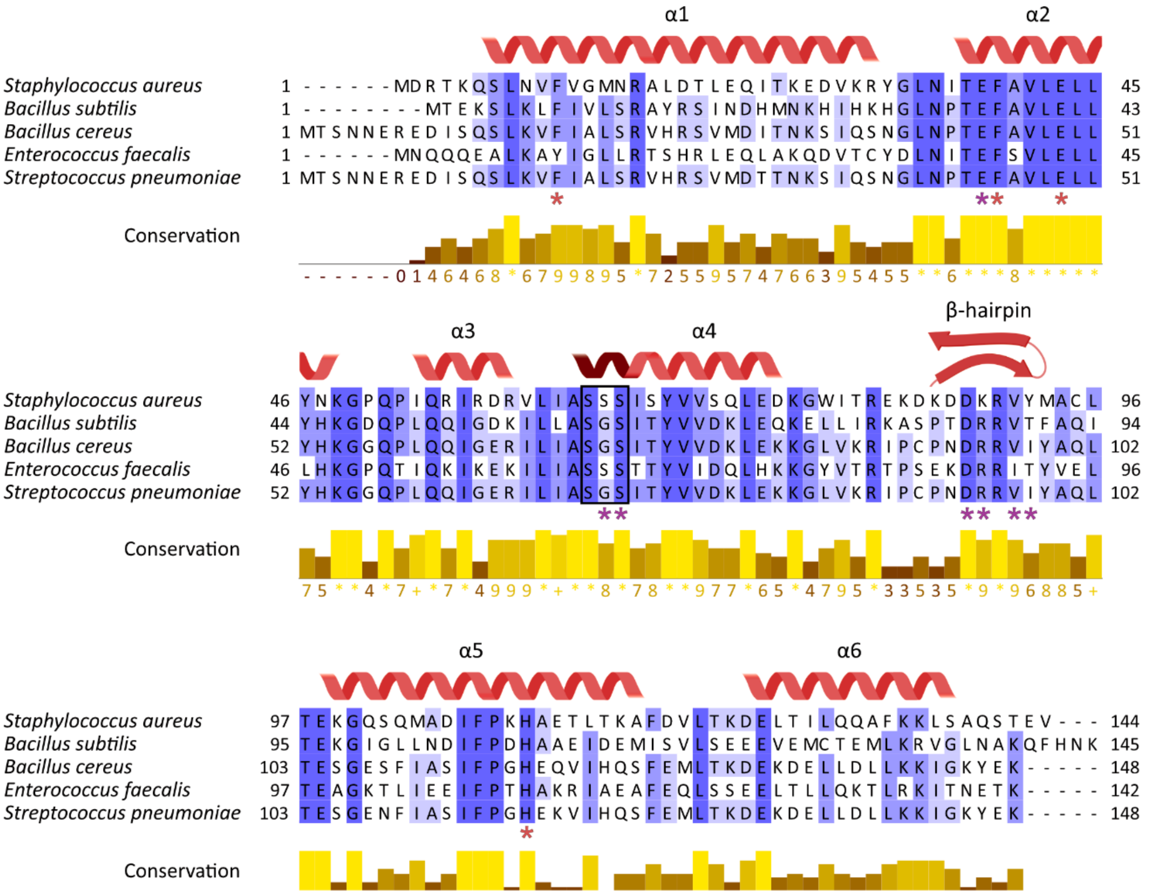


**Figure S14. Alignment of the MBQ-binding pocket and the allosteric loop region between MhqR homologs of Firmicutes.** Sequence alignment was performed with ClustalΩ2 for selected MhqR homologs of Firmicutes, including *Sa*MhqR, MhqR (BSU13670) of *B. subtilis*, CN491_21485 of *Bacillus cereus*, EF_1668 of *Enterococcus faecalis* and YusO of *Streptococcus pneumoniae.* The alignment supports the high conservation of the residues, forming the MBQ-binding pocket (F11, F39A, E43 and H111), the helix α2, and the α3-α4 allosteric loop region (LIASSSI) involved in redox regulation of DNA-binding activity of *Sa*MhqR. The MBQ pocket residues F11, F39A, E43 and H111 are marked by red stars. The serine residues (S65, S66) responsible for the elongation of α4 upon DNA interaction and residues in α2 (E38) and in the ß-hairpin (D88, K89, V91 and Y92), which are important for DNA-binding of *Sa*MhqR are marked by violet stars.

**Table S1.** Crystallographic data collection and refinement statistics for MhqR structures.

| | ***Sa*MhqR (apo)** | ***Sa*MhqR (MBQ-bound)** |
|---|---|---|
| **Data collection** | | |
| Space group | $P2_12_12_1$ | $P2_12_12_1$ |
| Cell dimensions | | |
| *a*, *b*, *c* (Å) | 51.92, 69.43, 71.65 | 47.34, 73.64, 95.16 |
| α, β, γ (°) | 90, 90, 90 | 90, 90, 90 |
| Resolution (Å) | 49.86 - 1.55 (1.59 - 1.55) | 47.58 - 2.28 (2.42 - 2.28) |
| $I/\sigma(I)$ | 11.04 (1.79) | 7.90 (2.68) |
| Completeness (%) | 99.8 (96.7) | 99.9 (99.9) |
| Redundancy | 13.0 (12.2) | 12.37 (11.68) |
| $CC_{1/2}$ | 0.998 (0.612) | 0.994 (0.918) |
| **Refinement** | | |
| Resolution (Å) | 49.86 - 1.55 (1.59 - 1.55) | 47.58 - 2.28 (2.42 - 2.28) |
| No. reflections | 38185 (2589) | 15743 (2484) |
| $R_{work}$ / $R_{free}$ | 0.200 (0.252) / 0.235 (0.272) | 0.240 (0.273) / 0.283 (0.349) |
| No. atoms | | |
| Protein | 2337 | 2318 |
| Ligand/ion | 20 | 54 |
| Water | 196 | 79 |
| Mean *B*-factors | | |
| Protein | 27.51 | 30.21 |
| Ligand/ion | 38.62 | 27.79 |
| Water | 33.87 | 30.33 |
| Ramachandran (%) | | |
| favored | 99.29 | 98.93 |
| allowed | 0.71 | 1.07 |
| outliers | 0.00 | 0.00 |
| r.m.s. deviations | | |
| Bond lengths (Å) | 0.015 | 0.008 |
| Bond angles (°) | 1.33 | 0.98 |

Values in parentheses denote the highest resolution shell.

**Table S2.** Thermodynamic values determined from isothermal titration calorimetry (ITC) experiments.

| **Experiment** | | | **$K_D$ (µM)** | **averaged $K_D$ (µM)** | **ΔH (kcal mol$^{-1}$)** | **-TΔS (kcal mol$^{-1}$)** | **ΔG (kcal)** | **N (sites)** |
|---|---|---|---|---|---|---|---|---|
| WT vs. | MHQ | R1: | 8.56 ± 0.586 | 7.09 ± 0.37 | -6.1 | -0.8 | -6.91 | 1.05 |
| | | R2: | 6.14 ± 0.472 | | -5.7 | -1.4 | -7.11 | 1.45 |
| | MBQ | R1: | 3.12 ± 0.613 | 3.39 ± 0.53 | -30.9 | 23.4 | -7.51 | 1.00 |
| | | R2: | 4.20 ± 1.070 | | -30.4 | 23.0 | -7.34 | 1.06 |
| F11A vs. | MHQ | R1: | 2.78 ± 0.249 | 2.05 ± 0.12 | -4.0 | -3.6 | -7.58 | 2.98 |
| | | R2: | 1.87 ± 0.146 | | -3.4 | -4.4 | -7.82 | 4 |
| | MBQ | R1: | 4.26 ± 0.812 | 4.32 ± 0.60 | -31.2 | 23.8 | -7.33 | 1.05 |
| | | R2: | 4.39 ± 0.877 | | -31.6 | 24.3 | -7.31 | 1.05 |
| H111A vs. | MHQ | R1: | 4.46 ± 0.295 | 4.67 ± 0.26 | -6.3 | -1.0 | -7.30 | 3.07 |
| | | R2: | 5.45 ± 0.548 | | -1.5 | -5.7 | -7.18 | 1* |
| | MBQ | R1: | 4.02 ± 0.815 | 4.23 ± 0.60 | -33.0 | 25.7 | -7.36 | 1.04 |
| | | R2: | 4.49 ± 0.904 | | -33.4 | 26.1 | -7.30 | 1.04 |
| E43A vs. | MHQ | R1: | 6.09 ± 0.407 | 5.88 ± 0.36 | -12.9 | 5.8 | -7.12 | 3.24 |
| | | R2: | 5.48 ± 0.826 | | -1.0 | -7.1 | -7.18 | 1* |
| | MBQ | R1: | 5.09 ± 0.803 | 5.32 ± 0.70 | -33.2 | 26.0 | -7.22 | 1.09 |
| | | R2: | 6.57 ± 1.710 | | -35.8 | 28.7 | -7.07 | 1.08 |
| F39A vs. | MHQ | R1: | 4.25 ± 0.311 | 4.35 ± 0.28 | -13.5 | 6.2 | -7.33 | 2.68 |
| | | R2: | 4.68 ± 0.857 | | -1.2 | -6.1 | -7.27 | 2.74 |
| | MBQ | R1: | 3.26 ± 0.344 | 2.90 ± 0.27 | -28.1 | 20.8 | -7.49 | 1.08 |
| | | R2: | 2.54 ± 0.441 | | -26.9 | 19.3 | -7.64 | 1.02 |

*An N (sites) of 1 was assigned to enable the curve fitting.

Averaged $K_D$ (µM) was calculated as weighted mean with propagated uncertainty.

**Table S3.** Bacterial strains, phage and plasmids.

| Strain | Description | Reference |
|---|---|---|
| ***Staphylococcus aureus*** | | |
| RN4220 | restriction negative strain/MSSA cloning intermediate derived from 8325-4 | [1] |
| COL | archaic HA-MRSA strain | [2] |
| COL-Δ*mhqR* | COL Δ*mhqR* deletion mutant | [3] |
| COL-Δ*mhqR*-pRB473-*mhqR* | COL Δ*mhqR* complemented with pRB473-*mhqR* | [3] |
| COL-Δ*mhqR*-pRB473-*mhqR* F11A | COL Δ*mhqR* complemented with pRB473-*mhqR* F11A | This study |
| COL-Δ*mhqR*-pRB473-*mhqR* E38A | COL Δ*mhqR* complemented with pRB473-*mhqR* E38A | This study |
| COL-Δ*mhqR*-pRB473-*mhqR* F39A | COL Δ*mhqR* complemented with pRB473-*mhqR* F39A | This study |
| COL-Δ*mhqR*-pRB473-*mhqR* E43A | COL Δ*mhqR* complemented with pRB473-*mhqR* E43A | This study |
| COL-Δ*mhqR*-pRB473-*mhqR* Q54A | COL Δ*mhqR* complemented with pRB473-*mhqR* Q54A | This study |
| COL-Δ*mhqR*-pRB473-*mhqR* A63E | COL Δ*mhqR* complemented with pRB473-*mhqR* A63E | This study |
| COL-Δ*mhqR*-pRB473-*mhqR* S64A | COL Δ*mhqR* complemented with pRB473-*mhqR* S64A | This study |
| COL-Δ*mhqR*-pRB473-*mhqR* S65A | COL Δ*mhqR* complemented with pRB473-*mhqR* S65A | This study |
| COL-Δ*mhqR*-pRB473-*mhqR* S66A | COL Δ*mhqR* complemented with pRB473-*mhqR* S66A | This study |
| COL-Δ*mhqR*-pRB473-*mhqR* S68A | COL Δ*mhqR* complemented with pRB473-*mhqR* S68A | This study |
| COL-Δ*mhqR*-pRB473-*mhqR* Y69A | COL Δ*mhqR* complemented with pRB473-*mhqR* Y69A | This study |
| COL-Δ*mhqR*-pRB473-*mhqR* V70A | COL Δ*mhqR* complemented with pRB473-*mhqR* V70A | This study |
| COL-Δ*mhqR*-pRB473-*mhqR* Q73A | COL Δ*mhqR* complemented with pRB473-*mhqR* Q73A | This study |
| COL-Δ*mhqR*-pRB473-*mhqR* D88A | COL Δ*mhqR* complemented with pRB473-*mhqR* D88A | This study |
| COL-Δ*mhqR*-pRB473-*mhqR* K89A | COL Δ*mhqR* complemented with pRB473-*mhqR* K89A | This study |
| COL-Δ*mhqR*-pRB473-*mhqR* R90A | COL Δ*mhqR* complemented with pRB473-*mhqR* R90A | This study |
| COL-Δ*mhqR*-pRB473-*mhqR* V91A | COL Δ*mhqR* complemented with pRB473-*mhqR* V91A | This study |
| COL-Δ*mhqR*-pRB473-*mhqR* Y92A | COL Δ*mhqR* complemented with pRB473-*mhqR* Y92A | This study |
| COL-Δ*mhqR*-pRB473-*mhqR* H111A | COL Δ*mhqR* complemented with pRB473-*mhqR* H111A | This study |
| COL-Δ*mhqR*-pRB473-*mhqR* E43A,H111A | COL Δ*mhqR* complemented with pRB473-*mhqR* E43A,H111A | This study |
| *Staphylococcus* phage 81 | | [4] |
| ***Escherichia coli*** | | |
| DH5α | F-φ80dlacZ Δ(lacZYA-argF) U169 deoRsupE44ΔlacU169 (f80lacZDM15) hsdR17 recA1 endA1 (rk- mk+) supE44gyrA96 thi-1 gyrA69 relA1 | [5] |
| BL21(DE3)*plysS* | F- ompT hsdS gal (rb- mb+) DE3(Sam7 Δnin5 lacUV5-T7 Gen1) | [5] |
| BL21(DE3)*plysS* pET11b-*mhqR* | For overexpression of His-tagged MhqR wild type | [3] |
| BL21(DE3)*plysS* pET11b-*mhqR* F11A | For overexpression of His-tagged MhqR F11A | This study |

| | | |
|---|---|---|
| BL21(DE3)*plysS* pET11b-*mhqR* E38A | For overexpression of His-tagged MhqR E38A | This study |
| BL21(DE3)*plysS* pET11b-*mhqR* F39A | For overexpression of His-tagged MhqR F39A | This study |
| BL21(DE3)*plysS* pET11b-*mhqR* E43A | For overexpression of His-tagged MhqR E43A | This study |
| BL21(DE3)*plysS* pET11b-*mhqR* Q54A | For overexpression of His-tagged MhqR Q54A | This study |
| BL21(DE3)*plysS* pET11b-*mhqR* A63E | For overexpression of His-tagged MhqR A63E | This study |
| BL21(DE3)*plysS* pET11b-*mhqR* S64A | For overexpression of His-tagged MhqR S64A | This study |
| BL21(DE3)plysS pET11b-*mhqR* S65A | For overexpression of His-tagged MhqR S65A | This study |
| BL21(DE3)plysS pET11b-*mhqR* S66A | For overexpression of His-tagged MhqR S66A | This study |
| BL21(DE3)plysS pET11b-*mhqR* S68A | For overexpression of His-tagged MhqR S68A | This study |
| BL21(DE3)plysS pET11b-*mhqR* Y69A | For overexpression of His-tagged MhqR Y69A | This study |
| BL21(DE3)plysS pET11b-*mhqR* V70A | For overexpression of His-tagged MhqR V70A | This study |
| BL21(DE3)plysS pET11b-*mhqR* Q73A | For overexpression of His-tagged MhqR Q73A | This study |
| BL21(DE3)plysS pET11b-*mhqR* D88A | For overexpression of His-tagged MhqR D88A | This study |
| BL21(DE3)plysS pET11b-*mhqR* K89A | For overexpression of His-tagged MhqR K89A | This study |
| BL21(DE3)plysS pET11b-*mhqR* R90A | For overexpression of His-tagged MhqR R90A | This study |
| BL21(DE3)plysS pET11b-*mhqR* V91A | For overexpression of His-tagged MhqR V91A | This study |
| BL21(DE3)plysS pET11b-*mhqR* Y92A | For overexpression of His-tagged MhqR Y92A | This study |
| BL21(DE3)*plysS* pET11b-*mhqR* H111A | For overexpression of His-tagged MhqR H111A | This study |
| BL21(DE3)*plysS* pET11b-*mhqR* E43A,H111A | For overexpression of His-tagged MhqR E43A,H111A | This study |
| **Plasmids** | | |
| pRB473 | pRB373-derivative, *E. coli*/ *S. aureus* shuttle vector, containing xylose-inducible $P_{Xyl}$ promoter $Amp^R$, $Cm^R$ | [6, 7] |
| pRB473-*mhqR* F11A | pRB473 expressing *mhqR* F11A under $P_{Xyl}$ | This study |
| pRB473-*mhqR* E38A | pRB473 expressing *mhqR* E38A under $P_{Xyl}$ | This study |
| pRB473-*mhqR* F39A | pRB473 expressing *mhqR* F39A under $P_{Xyl}$ | This study |
| pRB473-*mhqR* E43A | pRB473 expressing *mhqR* E43A under $P_{Xyl}$ | This study |
| pRB473-*mhqR* Q54A | pRB473 expressing *mhqR* Q54A under $P_{Xyl}$ | This study |
| pRB473-*mhqR* A63E | pRB473 expressing *mhqR* A63E under $P_{Xyl}$ | This study |
| pRB473-*mhqR* S64A | pRB473 expressing *mhqR* S64A under $P_{Xyl}$ | This study |
| pRB473-*mhqR* S65A | pRB473 expressing *mhqR* S65A under $P_{Xyl}$ | This study |
| pRB473-*mhqR* S66A | pRB473 expressing *mhqR* S66A under $P_{Xyl}$ | This study |
| pRB473-*mhqR* S68A | pRB473 expressing *mhqR* S68A under $P_{Xyl}$ | This study |
| pRB473-*mhqR* Y69A | pRB473 expressing *mhqR* Y69A under $P_{Xyl}$ | This study |
| pRB473-*mhqR* V70A | pRB473 expressing *mhqR* V70A under $P_{Xyl}$ | This study |
| pRB473-*mhqR* Q73A | pRB473 expressing *mhqR* Q73A under $P_{Xyl}$ | This study |
| pRB473-*mhqR* D88A | pRB473 expressing *mhqR* D88A under $P_{Xyl}$ | This study |
| pRB473-*mhqR* K89A | pRB473 expressing *mhqR* K89A under $P_{Xyl}$ | This study |
| pRB473-*mhqR* R90A | pRB473 expressing *mhqR* R90A under $P_{Xyl}$ | This study |
| pRB473-*mhqR* V91A | pRB473 expressing *mhqR* V91A under $P_{Xyl}$ | This study |
| pRB473-*mhqR* Y92A | pRB473 expressing *mhqR* Y92A under $P_{Xyl}$ | This study |
| pRB473-*mhqR* H111A | pRB473 expressing *mhqR* H111A under $P_{Xyl}$ | This study |
| pRB473-*mhqR* E43A,H111A | pRB473 expressing *mhqR* E43A,H111K under $P_{Xyl}$ | This study |

$^R$: resistant, Amp: ampicillin, Cm: chloramphenicol

**Table S4.** Oligonucleotide (primer) sequences

| Primer name | Sequence (5' to 3') | Reference |
|---|---|---|
| mhqR-pET- For-NheI | CTA<u>GCTAGC</u>ATGGATCGAACGAAACAATCTC | [3] |
| mhqR-pET-Rev-BamHI | CGC<u>GGATCC</u>TTAGTGATGGTGATGGTGATGCACTTCT GTAGATTGTGCACTT | [3] |
| F11A-for | CTA<u>GCTAGC</u>ATGGATCGAACGAAACAATCTCTCAATG TT**GCT**GTC | |
| E38A-for | GCGATATGGCTTAAATATTACTG**C**ATTTGCAG | |
| E38A-rev | AGCAACTCGAGCACTGCAAAT**G**CAGTAATATT | |
| F39A-for | AATATTACTGAA**GCT**GCAGTGCTCGAG | |
| F39A-rev | CTCGAGCACTGC**AGC**TTCAGTAATATT | |
| E43A-for | TTTGCAGTGCTCG**C**GTTGCTTTATAAT | |
| E43A-rev | ATTATAAAGCAAC**G**CGAGCACTGCAAA | |
| Q54A-for | TAATAAAGGTCCGCAACCAATT**GC**ACGTATTAGAG | |
| Q54A-rev | AATACGCGGTCTCTAATACGT**GC**AATTGGTTGCG | |
| A63E-for | CGCGTATTAATTG**A**AAGTAGCAGCATT | |
| A63E-rev | AATGCTGCTACTT**T**CAATTAATACGCG | |
| S64A-for | AGAGACCGCGTATTAATTGCA**GC**TAGCAGCATTTC | |
| S64A-rev | ACAACATATGAAATGCTGCTA**GC**TGCAATTAATAC | |
| S65A-for | TTAATTGCAAGT**GC**CAGCATTTCATAT | |
| S65A-rev | ATATGAAATGCTG**GC**ACTTGCAATTAA | |
| S66A-for | GCGTATTAATTGCAAGTAGC**GC**CATTTCATATGTT | |
| S66A-rev | TGACTTACAACATATGAAATG**GC**GCTACTTGCAAT | |
| S68A-for | TAATTGCAAGTAGCAGCATT**G**CATATGTTGTAAG | |
| S68A-rev | CCTCTAATTGACTTACAACATATG**C**AATGCTGCTAC | |
| Y69A-for | AATTGCAAGTAGCAGCATTTCA**GC**TGTTGTAAGTCA | |
| Y69A-rev | GTCCTCTAATTGACTTACAACA**GC**TGAAATGCTGC | |
| V70A-for | TGCAAGTAGCAGCATTTCATATG**C**TGTAAGTCA | |
| V70A-rev | CCTTTGTCCTCTAATTGACTTACA**G**CATATGAAATG | |
| Q73A-for | AGCATTTCATATGTTGTAAGT**GC**ATTAGAGGACA | |
| Q73A-rev | AACCTTTGTCCTCTAAT**GC**ACTTACAACATATG | |
| D88A-for | ACGTGAAAAGGATAAAGATG**C**TAAACGTGTATA | |
| D88A-rev | AAGCCATATATACACGTTTA**G**CATCTTTATCCTT | |
| K89A-for | TGAAAAGGATAAAGATGAT**GC**ACGTGTATATAT | |
| K89A-rev | TAAACAAGCCATATATACACGT**GC**ATCATCTTTAT | |
| R90A-for | GAAAAGGATAAAGATGATAAA**GC**TGTATATATGGC | |
| R90A-rev | CAGTTAAACAAGCCATATATACA**GC**TTTATCATCTT | |
| V91A-for | ATAAAGATGATAAACGTG**C**ATATATGGCTTG | |
| V91A-rev | TCAGTTAAACAAGCCATATAT**G**CACGTTTATCA | |
| Y92A-for | ATAAAGATGATAAACGTGTA**GC**TATGGCTTGTT | |
| Y92A-rev | TTCAGTTAAACAAGCCATA**GC**TACACGTTTATCA | |
| H111A-for | TTCCCTAAG**GCT**GCTGAGACATTAACAAAA | |
| H111A-rev | TTTTGTTAATGTCTCAGC**AGC**CTTAGGGAA | |
| qPCR-mhqE-for | TGCCTAACGATGACGCATTAAC | |
| qPCR-mhqE-rev | TGGCCATCGACTTCTTCAAATG | |
| mhqR-pRB-for-BamHI | TAG<u>GGATCC</u>TAAAATAAAAAAGTTGGTGATCATATGG ATCGAACGAAACAAT | [3] |
| mhqR-pRB-rev-KpnI | CTC<u>GGTACC</u>TTACACTTCTGTAGATTGTGCACTT | [3] |
| NB-mhqE-for | ACAGAAGTACTAGGCATGCGT | |
| NB-mhqE-rev | CTAATACGACTCACTATAGGGAGAACCCTTCGCCTAA TGTTTCA | |
| NB-mhqR-for | GGATCGAACGAAACAATCTCTCA | |
| NB-mhqR-rev | CTAATACGACTCACTATAGGGAGACGCTTTTGTTAAT GTCTCAGCA | |
| emsa2531-for | ACATATCAACTCCTATCATGATT | [3] |

| | | |
|---|---|---|
| emsa2531-rev | CGTTCGATCCATATGATCACC | [3] |

Restriction sites are underlined and bold bases indicate point mutations
